# Mechanisms underlying divergent relationships between Ca^2+^ and YAP/TAZ signaling

**DOI:** 10.1101/2022.10.06.511161

**Authors:** A. Khalilimeybodi, S.I. Fraley, P. Rangamani

**Affiliations:** Department of Mechanical and Aerospace Engineering, Jacobs School of Engineering, University of California San Diego, La Jolla CA 92093; Department of Bioengineering, Jacobs School of Engineering, University of California San Diego, La Jolla CA 92093

## Abstract

Yes-associated protein (YAP) and its homolog TAZ are transducers of several biochemical and biomechanical signals, serving to integrate multiplexed inputs from the microenvironment into higher-level cellular functions such as proliferation, differentiation, apoptosis, migration, and hemostasis. Emerging evidence suggests that Ca^2+^ is a key second messenger that closely connects microenvironmental input signals and YAP/TAZ regulation. However, studies that directly modulate Ca^2+^ have reported contradictory YAP/TAZ responses: In some studies, a reduction in Ca^2+^ influx increases the activity of YAP/TAZ, while in others, an increase in Ca^2+^ influx activates YAP/TAZ. Importantly, Ca^2+^ and YAP/TAZ exhibit distinct spatiotemporal dynamics, making it difficult to unravel their connections from a purely experimental approach. In this study, we developed a network model of Ca^2+^-mediated YAP/TAZ signaling to investigate how temporal dynamics and crosstalk of signaling pathways interacting with Ca^2+^ can alter YAP/TAZ response, as observed in experiments. By including six signaling modules (e.g., GPCR, IP3-Ca^2+^, Kinases, RhoA, F-actin, and Hippo-YAP/TAZ) that interact with Ca^2+^, we investigated both transient and steady-state cell response to Angiotensin II and thapsigargin stimuli. The model predicts stimuli, Ca^2+^ transient, and frequency-dependent relationships between Ca^2+^ and YAP/TAZ primarily mediated by signaling species like cPKC, DAG, CaMKII, and F-actin. Model results illustrate the role of Ca^2+^ dynamics and CaMKII bistable response in switching the direction of changes in Ca^2+^-induced YAP/TAZ activity for different stimuli. Frequency-dependent YAP/TAZ response revealed the competition between upstream regulators of LATS1/2, leading to the YAP/TAZ non-monotonic response to periodic GPCR stimulation. This study provides new insights into the underlying mechanisms responsible for the controversial Ca^2+^-YAP/TAZ relationship observed in experiments.

## 1 Introduction

In any given cell, numerous external biochemical and biomechanical cues with different dynamics act together to control the cell’s functions and phenotypic changes^1,2^. The intracellular interaction between these stimuli is orchestrated through a myriad signaling pathways enabling the cell to respond to various inputs proportional to their dynamics^3,4^. Yes-associated protein (YAP) and its homolog, transcriptional coactivator with PDZ-binding motif (TAZ), have recently attracted researchers’ attention as key targets for integrating biochemical and biomechanical signals^5,6^. The role of YAP/TAZ in regulating many cellular functions such as proliferation, differentiation, apoptosis, migration, and homeostasis has been demonstrated by the previous studies^7,8^. YAP/TAZ responds to different cues by shuttling between the cytoplasm (inactive state) and nucleus (active state) to activate its cofactors, such as TEA domain transcription factors (TEADs), and thereby regulate the cell functions^6,9^. Although the upstream pathways of YAP/TAZ activation have been explored^8,10^, the YAP/TAZ response when multiple upstream signals exhibiting different dynamics act together like in GPCR signaling is not straightforward^11^. This complexity and, in some cases, uncertainty in YAP/TAZ response (e.g., Ca^2+^-induced YAP/TAZ) prevents us from understanding the dynamics of cellular functions controlled by YAP/TAZ activity^12–14^.

The Hippo core cascade and the Rho family of GTPases (RhoA) pathways are primary transducers of several biochemical and biomechanical signals to YAP/TAZ^15^. The Hippo core cascade, including Mammalian STE20-like protein kinase 1/2 (MST1/2) and large tumor suppressors 1 and 2 (LATS1/2), is among the first discovered regulators of YAP/TAZ^16^, extensively studied in cancer research for its tumor suppressor function mainly through YAP/TAZ inhibition^17,18^. Numerous intrinsic and extrinsic signals can activate the Hippo pathway, including extracellular matrix (ECM) stiffness, cell polarity, cell-cell interaction, cellular energy status, and hormonal signals through GPCRs^19,20^. The RhoA pathway is an established and well-characterized master regulator of actin remodeling and cytoskeletal dynamics that can activate YAP/TAZ^21,22^. RhoA can be activated by mechanical stresses such as cell stretch, elevated ECM stiffness, and endogenous tension^23,24^. While the contribution of the Hippo and RhoA pathways to YAP/TAZ regulation is demonstrated by many studies^7,25,26^, there is a gap in the early upstream regulators of YAP/TAZ activity and their dynamics, especially by G-protein coupled receptors (GPCRs) agonists^11^. Ca^2+^ has recently been identified as a potential second messenger that closely connects the regulators of YAP/TAZ^27^ and actively participates in the YAP/TAZ regulation by mechanical and biochemical stimuli^28–30^. However, the contribution of Ca^2+^ to divergent responses of YAP/TAZ that have been observed in various experimental settings is not completely understood^27^. In this work, we aim to close this gap using computational modeling.

Ca^2+^ signaling can both inhibit and promote YAP/TAZ activity in various contexts, e.g., cell type, stimuli, and experiment conditions^27^ (Fig. 1A). In support of the inhibitory action of Ca^2+^ on YAP/TAZ, a study showed that Amlodipine, an L-type Ca^2+^ channel blocker, enhanced store-operated Ca^2+^ entry (SOCE) and led to elevated cytosolic Ca^2+^ level and the subsequent activation of the Hippo pathway resulting in the inhibition of YAP/TAZ^28^. Also, the knockout of two-pore channel 2 (TPC2) increased YAP/TAZ activity in metastatic melanoma cells by reducing Ca^2+^ influx^31^. On the other hand, some studies have demonstrated the activation of YAP/TAZ by Ca^2+^. For example, activation of the Piezo1 receptor by mechanical stimuli (ECM stiffness) increases Ca^2+^ influx through Piezo1 and results in YAP nuclear localization or activation in neural stem cells^32^. Another study found that cholesterol can activate TAZ by promoting the IP_3_R activity, leading to higher cytosolic Ca^2+ 29^. Part of this bidirectional response can be associated with the cell type-dependent response of YAP/TAZ to protein kinase C (PKC) isoforms^33^. Although all PKC isoforms have been demonstrated as regulators of YAP/TAZ activity in cells, their mechanisms of action are quite different. For example, while cPKCs promote YAP dephosphorylation (activation), nPKCs promote YAP phosphorylation, both through Hippo pathway regulation^33^. Moreover, the Ca^2+^-induced F-actin remodeling and subsequent elevation of cPKC (PKCβ) activity could lead to Hippo pathway activation and YAP/TAZ inhibition^27^. On the other hand, aPKC overexpression could increase YAP/TAZ activity by inhibiting MST1/2 from phosphorylating LATS1/2^34^. However, to our knowledge, no experimental or computational studies have investigated underlying mechanisms that drive the bidirectional changes in Ca^2+^-induced YAP/TAZ activity. Here, we investigate the contribution of PKC isoforms to Ca^2+^-induced YAP/TAZ activity.

**Figure 1.**
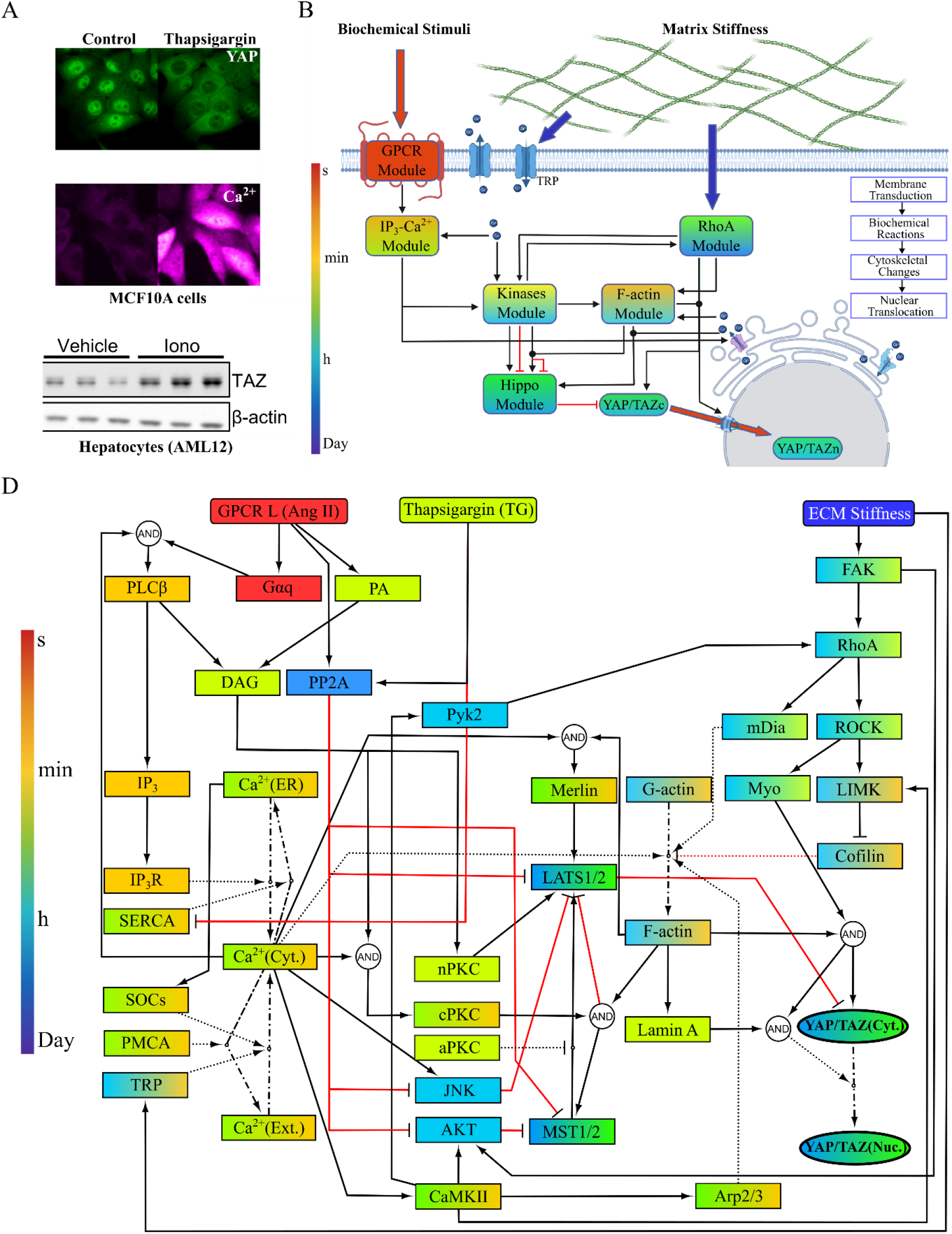
A compartmental model of Ca^2+^-mediated YAP/TAZ signaling to investigate the Ca^2+^-YAP/TAZ context-specific relationship. (A) Divergent Ca^2+^-YAP/TAZ relationships were observed in different experimental settings. An increase in the cytosolic Ca^2+^ level could both inhibit (MCF10A cells: adapted from ref.^56^, which is licensed under CC by 4.0.) or activate (AML12 cells: from ref.^29^, with permission from Elsevier) YAP/TAZ. (B) Signaling modules with different dynamics (time scales) that regulate YAP/TAZ activity after biomechanical and biochemical stimuli through Ca^2+^-mediated signaling are shown. IP_3_-Ca^2+^ module links fast GPCR module activation by biochemical stimuli, particularly angiotensin II (Ang II), to intracellular Ca^2+^ in a few minutes. RhoA module mediates the regulatory effect of slow biomechanical stimuli (ECM matrix stiffness) on YAP/TAZ activity from hours to days. Cell signaling modules, including F-actin, kinases, and Hippo, contribute to Ca^2+^-mediated YAP/TAZ regulation with various response times from minutes to hours. The time scale is shown in the color bar. (C) The network of signaling species and reactions connecting cell stimuli to YAP/TAZ activity. Thapsigargin (Tg), a SERCA inhibitor, is used in the model to modify intracellular Ca^2+^ directly. Activation and inhibition reactions are illustrated by black and red lines, respectively. Dot-dashed lines show species transfer between compartments. Dotted lines show species’ influence on reactions.

Our overarching hypothesis is that the different dynamics of signaling modules and species that link cell cues to Ca^2+^ and YAP/TAZ, as well as cell type-dependent differences in the Ca^2+^ transient, could modify Ca^2+^-YAP/TAZ interaction. We developed a dynamical systems model of Ca^2+^-mediated YAP/TAZ signaling based on prior knowledge to examine our hypothesis and calibrated our model with published experimental data. To investigate the effect of distinct biochemical signals on intracellular Ca^2+^ transients and subsequent YAP/TAZ regulation, we modeled Ca^2+^ dynamics induced by a Gq receptor agonist, Angiotensin II, and compared this to a SERCA inhibitor, thapsigargin. In the model, signal transduction from receptors to YAP/TAZ includes multiple biological processes such as membrane transduction, biochemical reactions, cytoskeletal changes, and nuclear translocation that take place on seconds-to-day timescales (Fig. 1B). The inherent difference in the dynamics of these processes enables the cells to produce long-term phenotypic changes in response to fast and transient input signals^35^. Building a model representing general non-excitable cells facilitates exploring the impact of different cell contexts, including cell stimuli (type and frequency) and cell type (Ca^2+^ transient), on YAP/TAZ activity through varying model parameters.

In summary, we mechanistically investigate the Ca^2+^-YAP/TAZ relationship and predict potential mechanisms that modify this relationship in the cell. We identified the main signaling species in Ca^2+^-mediated YAP/TAZ response and predicted how calcium/calmodulin-dependent protein kinase II (CaMKII) bistability could govern the switching from Ca^2+^-induced YAP/TAZ inhibition to activation and vice versa. By exploring the influence of Gq receptor stimulation frequency on YAP/TAZ activity, we predict a YAP/TAZ non-monotonic response to periodic GPCR stimulation mainly mediated by LATS1/2. Finally, the model can explain the diverse Ca^2+^-YAP/TAZ relationships reported in experimental studies.

## 2 Methods

### 2.1 Model development

Here, we describe the main features of our deterministic signaling model that investigates the crosstalk between intracellular Ca^2+^ dynamics and the activation of YAP/TAZ. We developed a compartmental ordinary differential equations (ODEs) model with seven (7) compartments to describe the dynamics of Ca^2+^-mediated YAP/TAZ activation. The seven compartments include four (4) volumetric compartments, including the extracellular matrix (ECM), cytosol (C), endoplasmic reticulum (ER), and nucleus (N), and three membranes, including plasma membrane (PM), ER membrane (ERM), and nuclear membrane (NM). The concentrations of species in volumetric compartments are given in μM. Membrane-associated species are represented by their density in the membrane (molecules/μm^2^). The model includes 74 signaling species linking biochemical and biomechanical inputs received by receptors on the plasma membrane to YAP/TAZ through 66 reactions. In the model, we assumed that all species are well mixed within their compartment. Interactions between compartments are represented through fluxes.

The list of model species, including their best guess initial values obtained from previous studies or assumed in this study, is presented in Table 1. We simulated cell response from a steady-state resting condition achieved by simulating the model without any inputs for a long time (100,000 s). These concentrations are provided in Table 1 and used as initial conditions for the main simulations in the result section. The reactions have been modeled mainly by mass action, Michaelis–Menten, and Hill kinetics originating from previously published models^36–39^. Tables 2-4 describe the reactions used in the model, including the reaction rate formula and kinetic parameters directly obtained from previous studies or estimated from experimental data. In tables, H(x) is equal to x for x>0 and zero for x<=0. We focused on a generic mammalian cell rather than a specific cell type in our model. Therefore, we used previously published computational models and empirical data from various non-excitable cell types to develop our model and estimate the parameters.

**Table 1.**
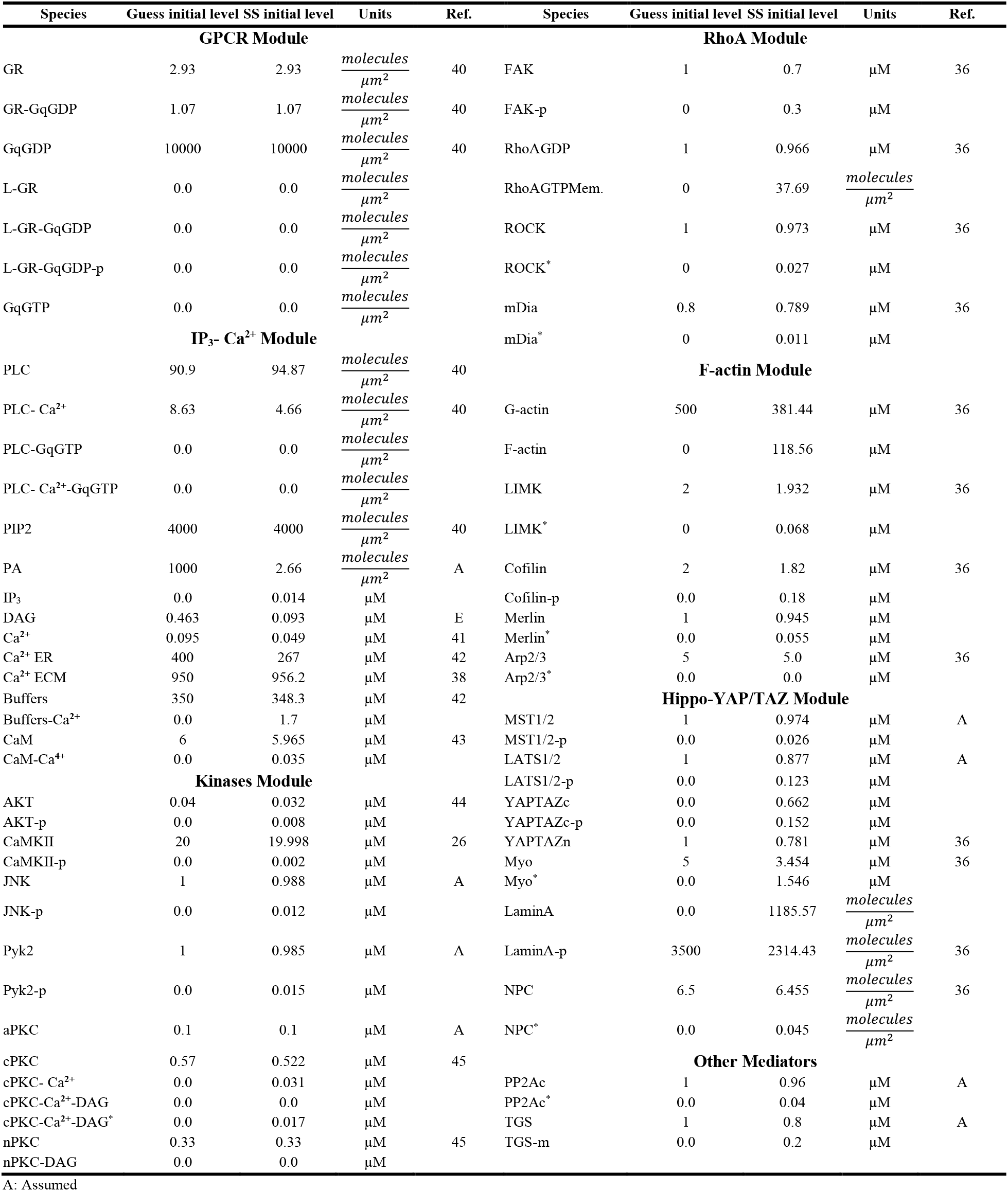
List of species in the Ca^2+^-mediated YAP/TAZ model

| Species | Guess initial level | SS initial level | Units | Ref. | Species | Guess initial level | SS initial level | Units | Ref. |
| --- | --- | --- | --- | --- | --- | --- | --- | --- | --- |
| **GPCR Module** |  |  |  |  | **RhoA Module** |  |  |  |  |
| GR | 2.93 | 2.93 | $\frac{\text{molecules}}{\mu\text{m}^2}$ | 40 | FAK | 1 | 0.7 | $\mu\text{M}$ | 36 |
| GR-GqGDP | 1.07 | 1.07 | $\frac{\text{molecules}}{\mu\text{m}^2}$ | 40 | FAK-p | 0 | 0.3 | $\mu\text{M}$ | |
| GqGDP | 10000 | 10000 | $\frac{\text{molecules}}{\mu\text{m}^2}$ | 40 | RhoAGDP | 1 | 0.966 | $\mu\text{M}$ | 36 |
| L-GR | 0.0 | 0.0 | $\frac{\text{molecules}}{\mu\text{m}^2}$ | | RhoAGTPMem. | 0 | 37.69 | $\frac{\text{molecules}}{\mu\text{m}^2}$ | |
| L-GR-GqGDP | 0.0 | 0.0 | $\frac{\text{molecules}}{\mu\text{m}^2}$ | | ROCK | 1 | 0.973 | $\mu\text{M}$ | 36 |
| L-GR-GqGDP-p | 0.0 | 0.0 | $\frac{\text{molecules}}{\mu\text{m}^2}$ | | ROCK* | 0 | 0.027 | $\mu\text{M}$ | |
| GqGTP | 0.0 | 0.0 | $\frac{\text{molecules}}{\mu\text{m}^2}$ | | mDia | 0.8 | 0.789 | $\mu\text{M}$ | 36 |
| **IP<sub>3</sub>-Ca<sup>2+</sup> Module** | | | | | mDia* | 0 | 0.011 | $\mu\text{M}$ | |
| PLC | 90.9 | 94.87 | $\frac{\text{molecules}}{\mu\text{m}^2}$ | 40 | **F-actin Module** | | | | |
| PLC-Ca<sup>2+</sup> | 8.63 | 4.66 | $\frac{\text{molecules}}{\mu\text{m}^2}$ | 40 | G-actin | 500 | 381.44 | $\mu\text{M}$ | 36 |
| PLC-GqGTP | 0.0 | 0.0 | $\frac{\text{molecules}}{\mu\text{m}^2}$ | | F-actin | 0 | 118.56 | $\mu\text{M}$ | |
| PLC-Ca<sup>2+</sup>-GqGTP | 0.0 | 0.0 | $\frac{\text{molecules}}{\mu\text{m}^2}$ | | LIMK | 2 | 1.932 | $\mu\text{M}$ | 36 |
| PIP2 | 4000 | 4000 | $\frac{\text{molecules}}{\mu\text{m}^2}$ | 40 | LIMK* | 0 | 0.068 | $\mu\text{M}$ | |
| PA | 1000 | 2.66 | $\frac{\text{molecules}}{\mu\text{m}^2}$ | A | Cofilin | 2 | 1.82 | $\mu\text{M}$ | 36 |
| IP<sub>3</sub> | 0.0 | 0.014 | $\mu\text{M}$ | | Cofilin-p | 0.0 | 0.18 | $\mu\text{M}$ | |
| DAG | 0.463 | 0.093 | $\mu\text{M}$ | E | Merlin | 1 | 0.945 | $\mu\text{M}$ | |
| Ca<sup>2+</sup> | 0.095 | 0.049 | $\mu\text{M}$ | 41 | Merlin* | 0.0 | 0.055 | $\mu\text{M}$ | |
| Ca<sup>2+</sup> ER | 400 | 267 | $\mu\text{M}$ | 42 | Arp2/3 | 5 | 5.0 | $\mu\text{M}$ | 36 |
| Ca<sup>2+</sup> ECM | 950 | 956.2 | $\mu\text{M}$ | 38 | Arp2/3* | 0.0 | 0.0 | $\mu\text{M}$ | |
| Buffers | 350 | 348.3 | $\mu\text{M}$ | 42 | **Hippo-YAP/TAZ Module** | | | | |
| Buffers-Ca<sup>2+</sup> | 0.0 | 1.7 | $\mu\text{M}$ | | MST1/2 | 1 | 0.974 | $\mu\text{M}$ | A |
| CaM | 6 | 5.965 | $\mu\text{M}$ | 43 | MST1/2-p | 0.0 | 0.026 | $\mu\text{M}$ | |
| CaM-Ca<sup>4+</sup> | 0.0 | 0.035 | $\mu\text{M}$ | | LATS1/2 | 1 | 0.877 | $\mu\text{M}$ | A |
| **Kinases Module** | | | | | LATS1/2-p | 0.0 | 0.123 | $\mu\text{M}$ | |
| AKT | 0.04 | 0.032 | $\mu\text{M}$ | 44 | YAPTAZc | 0.0 | 0.662 | $\mu\text{M}$ | |
| AKT-p | 0.0 | 0.008 | $\mu\text{M}$ | | YAPTAZc-p | 0.0 | 0.152 | $\mu\text{M}$ | |
| CaMKII | 20 | 19.998 | $\mu\text{M}$ | 26 | YAPTAZn | 1 | 0.781 | $\mu\text{M}$ | 36 |
| CaMKII-p | 0.0 | 0.002 | $\mu\text{M}$ | | Myo | 5 | 3.454 | $\mu\text{M}$ | 36 |
| JNK | 1 | 0.988 | $\mu\text{M}$ | A | Myo* | 0.0 | 1.546 | $\mu\text{M}$ | |
| JNK-p | 0.0 | 0.012 | $\mu\text{M}$ | | LaminA | 0.0 | 1185.57 | $\frac{\text{molecules}}{\mu\text{m}^2}$ | |
| Pyk2 | 1 | 0.985 | $\mu\text{M}$ | A | LaminA-p | 3500 | 2314.43 | $\frac{\text{molecules}}{\mu\text{m}^2}$ | 36 |
| Pyk2-p | 0.0 | 0.015 | $\mu\text{M}$ | | NPC | 6.5 | 6.455 | $\frac{\text{molecules}}{\mu\text{m}^2}$ | 36 |
| aPKC | 0.1 | 0.1 | $\mu\text{M}$ | A | NPC* | 0.0 | 0.045 | $\frac{\text{molecules}}{\mu\text{m}^2}$ | |
| cPKC | 0.57 | 0.522 | $\mu\text{M}$ | 45 | **Other Mediators** | | | | |
| cPKC-Ca<sup>2+</sup> | 0.0 | 0.031 | $\mu\text{M}$ | | PP2Ac | 1 | 0.96 | $\mu\text{M}$ | A |
| cPKC-Ca<sup>2+</sup>-DAG | 0.0 | 0.0 | $\mu\text{M}$ | | PP2Ac* | 0.0 | 0.04 | $\mu\text{M}$ | |
| cPKC-Ca<sup>2+</sup>-DAG* | 0.0 | 0.017 | $\mu\text{M}$ | | TGS | 1 | 0.8 | $\mu\text{M}$ | A |
| nPKC | 0.33 | 0.33 | $\mu\text{M}$ | 45 | TGS-m | 0.0 | 0.2 | $\mu\text{M}$ | |
| nPKC-DAG | 0.0 | 0.0 | $\mu\text{M}$ | | | | | | |

A: Assumed

**Table 2.**
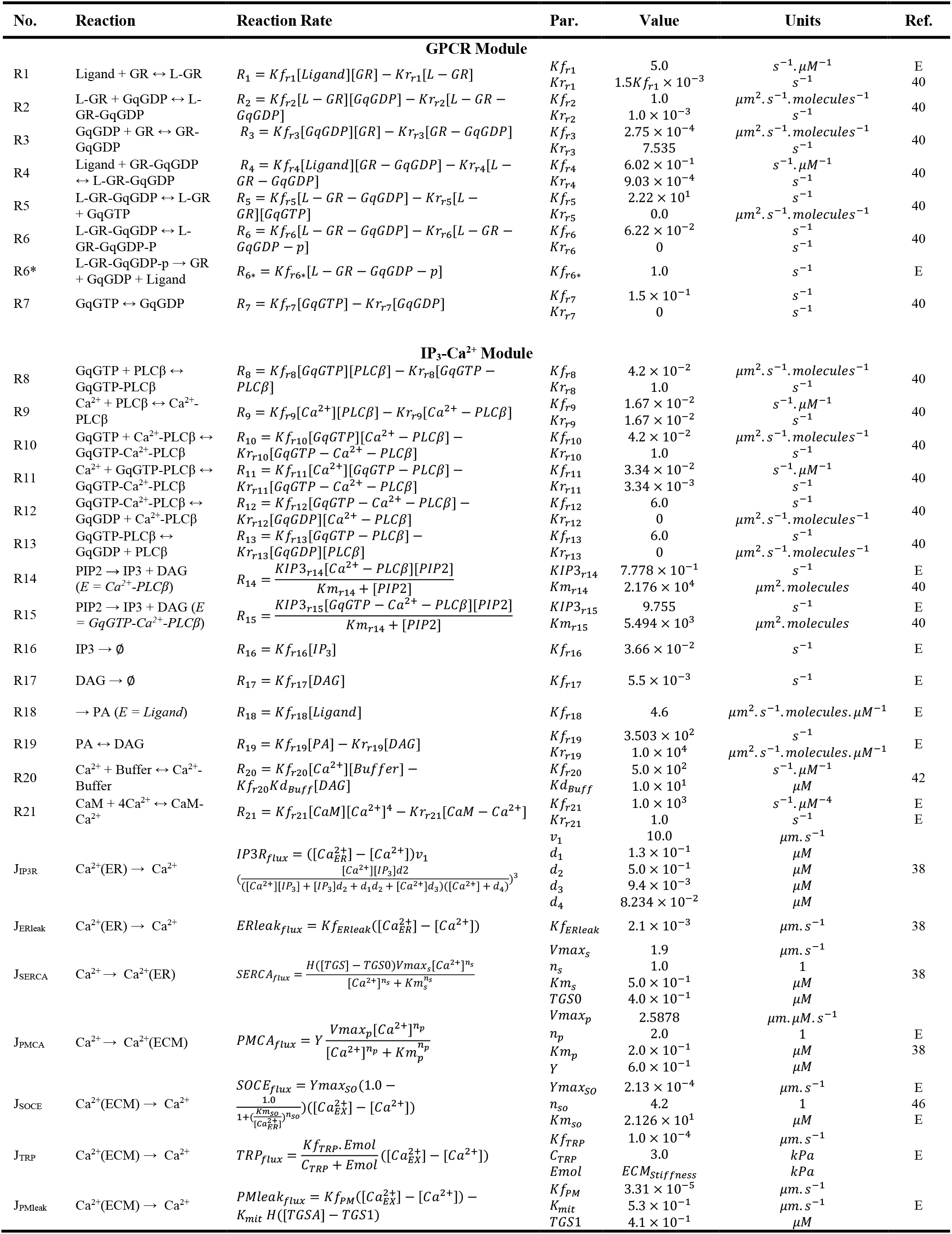
List of reactions and parameters in the GPCR and IP_3_-Ca^2+^ modules

**Table 3.**
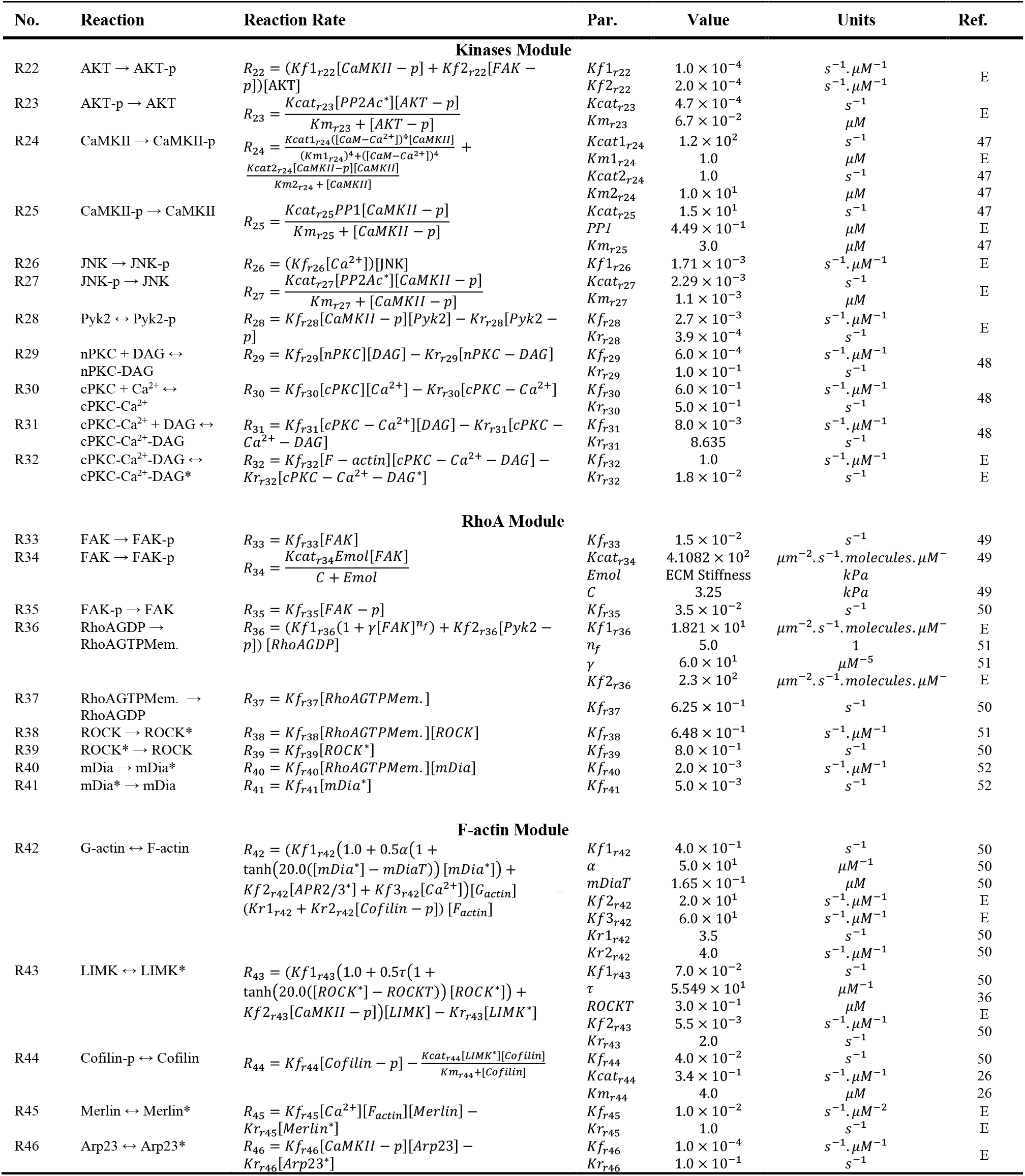
List of reactions and parameters in the Kinases, RhoA, and F-actin modules

**Table 4.**
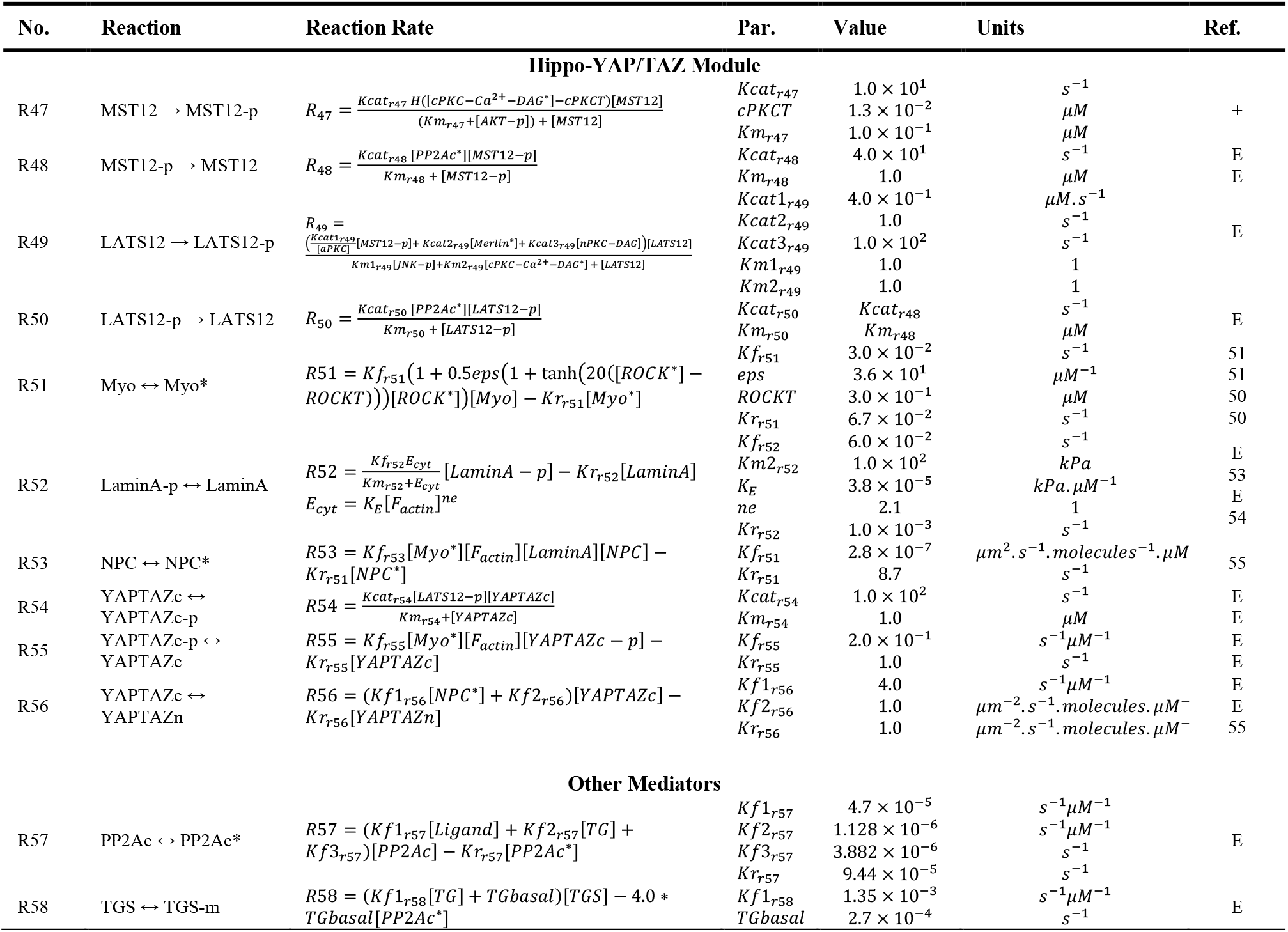
List of reactions and parameters in the Hippo-YAP/TAZ module

### 2.2 Main signaling modules

To understand the dynamics of Ca^2+^-mediated pathways and evaluate their contribution to YAP/TAZ activity, we developed a model involving main pathways that connect biochemical and biomechanical signals to YAP/TAZ through Ca^2+^. The model was constructed in a modular fashion to incorporate different pathways involved in Ca^2+^-mediated YAP/TAZ signaling and is illustrated in Fig.1B. The model involves six (6) closely interacting modules (e.g., GPCR, IP_3_-Ca^2+^, Kinases, RhoA, F-actin, and Hippo-YAP/TAZ) with Ca^2+^as the primary second messenger. As shown in Fig. 1B, the response times and dynamics of the modules cover a relatively broad range of timescales and behaviors due to the diversity in cell types, input signals, and signaling species in each module. For example, the GPCR module responds in a significantly shorter time to its inputs than the slower modules like the Hippo. Other modules/species with an intermediate response time, such as the kinases module, link these faster and slower modules. In the following, we discuss the main signaling modules contributing to the Ca^2+^-mediated YAP/TAZ regulation.

#### Module 1: GPCR activation

We used a mathematical model of Gq receptor cycling for non-excitable cells^40^. In this module, an extracellular ligand, angiotensin (Ang II), binds to cell-surface Gq receptor (GR) through reversible reactions causing a conformational change followed by the replacement of GDP with GTP on the Gqα subunit. Then, the Gqα subunit dissociates and stimulates the enzyme PLCβ. Phosphorylation of the active Gq receptor inactivates the receptor attenuating further signal transduction. To simulate cell response to Ang II, we modeled the ligand as a Heaviside step function with a nominal concentration of 0.1 µM. The default version of the GPCR module^40^ only becomes active for one cycle after ligand stimulation. For simulating cyclic activation of the GPCR module, we added reaction R6* (see Table 2) that resets the GPCR components to their initial states. The reaction R6* is only active during the OFF period of cyclic ligand activation. Reactions in the GPCR module were modeled by mass action kinetics (see Table 2).

#### Module 2: IP_3_-Ca^2+^ Module

In the IP_3_-Ca^2+^ module, the active Gqα subunit (GqGTP) binds through the reversible reactions to the phospholipase C β (PLCβ) and activates it. However, GqGTP can also become inactivated to GqGDP without any potential to activate PLCβ. In addition to GqGTP, calcium ions (Ca^2+^) are also necessary for the PLCβ activation (Fig. 1C). In the resting condition, PLCβ-Ca^2+^ hydrolyzes phosphatidylinositol 4,5-bisphosphate (PIP2) and generates inositol 1,4,5-trisphosphate (IP_3_) and diacylglycerol (DAG). However, after Gq stimulation with Ang II, PLCβ-Ca^2+^-GqGTP hydrolyzes PIP2 at a higher rate to produce IP_3_ and DAG. Furthermore, after Ang II stimulation, the activation of phospholipase D (PLD) leads to phosphatidic acid (PA) accumulation, which increases DAG concentration through its phosphohydrolase^57^. Thus, we included IP_3_ and DAG accumulation mechanisms by PIP2 hydrolysis through PLCβ (adopted from ref.^40^), Ang II-induced DAG accumulation by PA, and degradation of IP_3_ and DAG in our model.

In the model, Ca^2+^ levels in three ECM, cytosol, and ER compartments have been regulated by three fluxes through ER membrane, including J_IP3R_, J_SERCA_, and J_ERleak_, and four fluxes through the plasma membrane, including J_SOCE_, J_PMCA_, J_TRP_, and J_PMleak_. We did not include voltage-gated Ca^2+^ channels and ryanodine receptors (RyRs), because we focus on non-excitable cells^58^. In the model, J_IP3R_ represents the release of Ca^2+^ from ER to the cytosol after activation of the IP_3_R receptor by IP_3_^38,41^. We employed De Young and Keizer’s detailed model^59^ for the activation/deactivation of IP_3_R channels capturing the biphasic behavior of the IP_3_R channel opening at low and extremely high concentrations of Ca^2+^ and the CICR effect^37^. J_SERCA_ and J_PMCA_ are Ca^2+^ pump fluxes in the model (see Table 2)^38^.

J_ERleak_ and J_PMleak_ represent the Ca^2+^ leak from ER and plasma membranes, respectively, and primarily regulate the resting level of Ca^2+^ in the cell. Blocking SERCA by Tg disrupts Ca^2+^ homeostasis and causes the release of ER Ca^2+^ into the cytosol. Part of this Ca^2+^ enters mitochondria and results in mitochondrial fragmentation^60^. As mitochondria and their Ca^2+^ variations are not in the scope of this study, the Tg-induced Ca^2+^ efflux from the cytosol into mitochondria was modeled as part of the PM leak. J_SOCE_ and J_TRP_ are important routes for Ca^2+^ flux from ECM into the cytosol in non-excitable cells. In SOCE, the depletion of ER Ca^2+^ triggers translocation of STIM1 to Orai1 channels at PM and Ca^2+^ influx into the cytosol by activated Orai1 channels^61,62^. The overall process of Ca^2+^ depletion, STIM1 oligomerization, and translocation is a steep function of ER Ca^2+^ concentration, with a Hill coefficient in the range of 4 to 8^63^, and is included in our model by formulating J_SOCE_ as a Hill function of ER Ca^2+^. Furthermore, the transient receptor potential vanilloid-type 4 (TRPV4) channel is a matrix stiffness-sensitive Ca^2+^ channel that increases Ca^2+^ influx to cytosol in cells seeded on higher matrix stiffness^64^. We formulated J_TRP_ as a function of ECM stiffness.

#### Module 3: Kinases Module

Several kinases at Ca^2+^ downstream are known to mediate its effect on YAP/TAZ activity. As shown in Fig.1B, we identified the main players, including protein kinase C (PKC), CaMKII, protein kinase B (AKT), c-Jun N-terminal kinase (JNK), and protein tyrosine kinase 2 (Pyk2), from the literature that could contribute to YAP/TAZ regulation in a Ca^2+^-dependent manner. PKC represents one of the primary effectors downstream of GPCRs (especially Gq receptors) and Ca^2+^. PKC regulates a broad range of biological processes and can be classified into three sub-groups based on their activation mechanisms: 1) conventional PKCs (cPKC) that require both Ca^2+^ and DAG to be activated, 2) novel PKCs (nPKC) that are activated by only DAG, and 3) atypical PKCs (aPKC) that are independent of Ca^2+^ and DAG. CaMKII is another major kinase in Ca^2+^ signaling that regulates the activity of many species, including YAP/TAZ. As shown in Fig. 1C, CaMKII can control Actin Related Protein 2/3 complex (Arp2/3), LIM kinase (LIMK), AKT, and Pyk2 activity by phosphorylation. Arp2/3 and LIMK mediate CaMKII’s impact on F-actin remodeling by regulating the G-actin to F-actin transition^65,66^. AKT phosphorylates MST1/2 and inhibits its activity^67^, and PyK2 links Ca^2+^ and CaMKII activity to the RhoA pathway^68^. The c-Jun N-terminal kinase (JNK) also contributes to YAP activation by stiffness^69^. Activation of JNK by Ca^2+ 70^ can regulate Hippo pathway activity by phosphorylation of LATS1/2^25^.

#### Module 4: RhoA Module

In the RhoA module, focal adhesion kinase (FAK) links ECM stiffness and RhoA activation. An increase in ECM stiffness activates a cluster of integrins and their associated proteins, leading to FAK phosphorylation and subsequent activation of membrane-associated RhoA GTP^71^. Rho-associated kinase (ROCK) and mammalian diaphanous-related formin (mDia) are downstream targets of RhoA. ROCK mediates RhoA activity to the myosin light chain (Myo)^72^ as well as the F-actin module via LIMK^73^. Active mDia can facilitate F-actin polymerization^74^. For modeling the RhoA module, we employed a modified version of the YAP/TAZ signaling model^36^. We added Pyk2-dependent activation of RhoA and recalibrated certain parameters (Table 3) to capture ECM stiffness-induced variation of species in the RhoA module. We utilized steady-state data for the activity of species after an increase in ECM stiffness because of limited time-course data on ECM stiffness-induced variations in the biochemical activity of the species, which partly originates from the long time-scale of sensing of elevated stiffness by cells^75^.

#### Module 5: F-actin Module

In the F-actin module, cytoskeletal reorganization represented by the transition between G-actin to F-actin and vice versa is modeled. The F-actin module also includes LIMK, cofilin, Arp2/3, and moesin-ezrin-radixin like (Merlin), which govern interactions between F-actin dynamics and the activity of other signaling modules in the model. Activation of LIMK by ROCK or CaMKII phosphorylates cofilin and decreases its inhibitory effect on G-actin to F-actin transition^73^. Arp2/3 complex mediates the effect of CaMKII activation on F-actin polymerization^26,65^. The NF2 gene product, Merlin, is a cytoskeletal protein that regulates LATS1/2 activation. As shown by Wei et al.^76^, Ca^2+^ elevation leads to Merlin ubiquitination and then promotes LATS1/2 activation by facilitating the interaction between Merlin and LATS1/2. Rearranged F-actin due to Ca^2+^ elevation provides the scaffold for the Hippo pathway regulation by Merlin^27^. The F-actin module was developed by adding Arp2/3 and Merlin activation mechanisms via Mass Action kinetics to a previously developed YAP/TAZ signaling model^36^ (see Table 3).

#### Module 6: Hippo-YAP/TAZ Module

The Hippo signaling module comprises several kinases that target YAP/TAZ. MST1/2 and LATS1/2 are the main species in the Hippo pathway. Following the sequential activation of MST1/2 and LATS1/2 through their phosphorylation, active LATS1/2 phosphorylates YAP/TAZ, leading to the sequestration of YAP/TAZ in the cytoplasm^25^. If the Hippo pathway remains inactive, unphosphorylated YAP/TAZ translocates to the nucleus, where it binds to several cofactors, such as TEAD, that regulate the cell functions like growth and homeostasis through gene expression^8^. Furthermore, other mechanisms, such as the formation of stress fibers from myosin and F-actin, could facilitate the dephosphorylation of YAP/TAZ<u>^7^</u>.

Several studies^36,53,77^ demonstrated that species like LaminA and nuclear pore complexes (NPCs) could also regulate YAP/TAZ nuclear translocation. To model the LaminA and NPCs mechanisms of action, we utilized a previously published model^36^, which considers the LaminA dephosphorylation rate as a function of cell cytosolic stiffness. In that model, cytosolic stiffness is estimated as a function of F-actin. Then, LaminA, in coordination with other cytoskeletal proteins like F-actin and myosin, can lead to the elevated nuclear YAP/TAZ level by inducing nuclear stress and NPCs stretching resulting in lowered resistance of the NPC against the nuclear translocation of YAP/TAZ^36^.

#### Non-modular species

In addition to the signaling modules discussed above, our model has other signaling species to capture cellular responses in Ca^2+^-mediated YAP/TAZ activation. Protein Phosphatase 2A (PP2A) is a ubiquitously expressed serine-threonine phosphatase in cells that dephosphorylates many cellular species, including AKT, JNK, MST1/2, and LATS1/2^78,79^. Previous studies demonstrated elevated phosphatase activity of PP2A in response to Ang II and thapsigargin, the model inputs, in different cell types^80,81^. We used Michaelis–Menten kinetics to model the dephosphorylation of species by PP2A in our model. Furthermore, to model the inhibitory effect of Tg on SERCA, we defined a hypothetical species that acts as the substrate of Tg (TGS) and controls SERCA flux (J_SERCA_). TGS also regulates the cytosolic Ca^2+^ efflux to ECM to artificially reproduce the delayed decline in cytosolic Ca^2+^ level due to Ca^2+^ efflux into mitochondria after mitochondria fragmentation by Tg^60^.

### 2.3 Numerical methods

The model was constructed in VCell^82^. The system of deterministic ordinary differential equations was exported from VCell to MATLAB R2020b and was solved using the ‘ode15s’ solver for all numerical simulations with automatic time-stepping. The model was run for 100,000s without any inputs to reach the cell’s resting condition before conducting main simulations. We utilized a hybrid optimization algorithm comprising particle swarm optimization^83^ and a fmincon Trust Region Reflective Algorithm^84^ to obtain estimated parameters in MATLAB with higher accuracy. Steady-state values of model species were obtained at 100,000s when all species reached steady state. The model’s MATLAB code, including model species, parameters, reaction rates, and systems of ODEs, is available at https://github.com/mkm1712/Calcium_YAP-TAZ. Cell geometric parameters used for model simulation include cytosolic volume (2300 µm^3^), nuclear volume (550 µm^3^), ER volume (425 µm^3^), plasma membrane area (1260 µm^2^), nuclear membrane are (393 µm^2^), and ER membrane area (21000 µm^2^).

### 2.4 Model calibration to temporal and steady-state data

Out of the 156 kinetic parameters present in the model, 97 were obtained from previous models. To capture the dynamics of Ca^2+^-mediated YAPTAZ signaling, kinetic parameters that were not available directly from previous experimental or computational studies were estimated from available experimental data manually curated from the literature. We obtained either quantitative time course and steady-state data from research articles with similar cell types, assays, and experimental conditions with a preference for data from non-excitable endothelial cells. However, considering the limited data availability, a few sets of smooth muscle data were also included in the model calibration.

Figure 2 illustrates the comparison between model simulation results and experimental data for various model species. We utilized the scatter index (SI), which is the root mean square error (RMSE) normalized to the mean of measured data, to evaluate the goodness of fit between *in silico* results and calibration data. We conducted two types of calibrations – a comparison of the dynamics of concentrations of different species in the model for chemical stimuli (Fig. 2) and a comparison of the steady-state response of some species in the model as a function of substrate stiffness (Fig. S1). We show that the model can capture the dynamics of signaling species with different time scales after GPCR (Ang II) and thapsigargin (Tg) stimulations (Fig. 2). All experimental and *in silico* data were normalized to their maximum values to enable direct comparison. Figures 2A and B display simulated IP_3_ and cytosolic Ca^2+^ transient levels after Ang II stimulation reproducing experimental data with a peak in 20 seconds and returning to the resting level after 5 min. For DAG and PA, we observed a sustained increase in their concentrations after Ang II stimulation (Figs. 2C-D) with a steady-state level after 15 min. We also included Tg in the model to simulate direct regulation of Ca^2+^ to prevent potential interference of Ca^2+^-independent pathways in regulating YAP/TAZ response.

**Figure 2.**
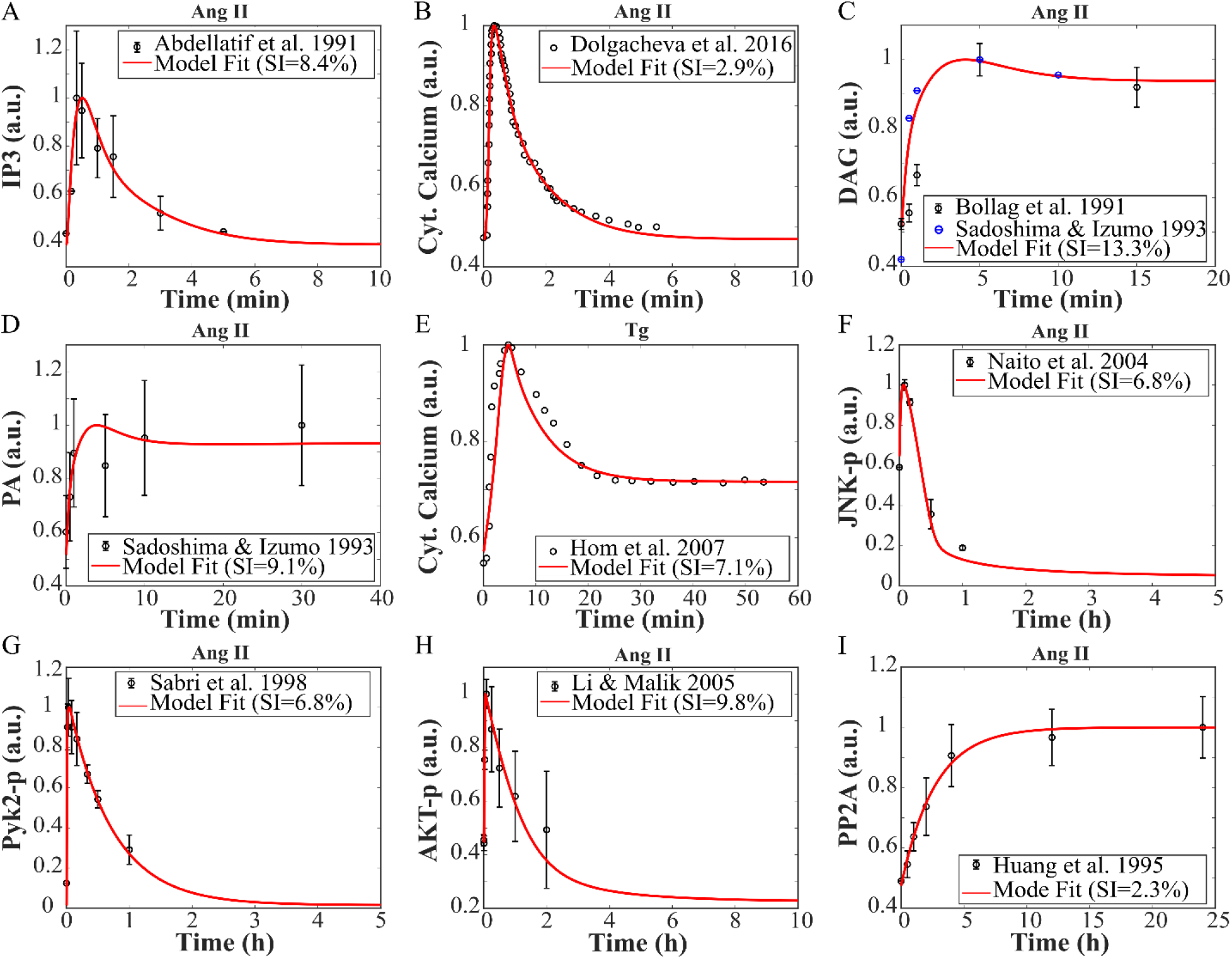
The compartmental ODE model captures temporal dynamics of Ca^2+^-mediated YAP/TAZ regulation in the cell. The model is calibrated to time course variations of IP_3_^87^(A), cytosolic Ca^2+ 88^ (B), DAG^57,89^ (C), PA^57^ (D), JNK^90^ (F), Pyk2^91^ (G), AKT^92^ (H), and PP2A^81^ (I) after Ang II (0.1μM) stimulation as well as variations of cytosolic Ca^2+^ after 1μM thapsigargin^60^ (E).

In contrast to Ang II, Tg-induced cytosolic Ca^2+^ transient in cells had a higher steady-state level than resting Ca^2+^ level due to the release of a significant amount of Ca^2+^ from ER to the cytosol (Fig. 2E). While a large portion of this Ca^2+^ is transported into mitochondria and ECM^60^, the remaining Ca^2+^ in cytosol leads to an elevated steady-state level of cytosolic Ca^2+^ after Tg stimulation^85,86^. Furthermore, Ang II induces a fast rising to the peak (2-5 min) in JNK, Pyk2, and AKT (Figs. 2F-H). However, their slow dynamics result in a long time (1-5 hours) for their concentrations to decline to baseline values. Interestingly, JNK and AKT have steady-state levels lower than their initial resting levels after Ang II stimulation, partly because of an Ang II-induced increase in PP2A phosphatase activity (Fig. 2I).

To develop a Ca^2+^-mediated YAP/TAZ signaling model capable of capturing various experimental settings, including potential changes in ECM stiffness, we utilized steady-state data of ECM stiffness as input to estimate some parameters in the RhoA and YAP/TAZ modules. Figures S1A and B illustrate the increase in the RhoAGTP^51^ and active myosin (Myo*)^51^ levels, respectively, in response to a higher ECM stiffness. RhoAGTP reaches its saturation level for ECM stiffness greater than 100 kPa, making its downstream target, including YAP/TAZ, insensitive to substrate (ECM) stiffness larger than 100 kPa. As suggested by some studies^36,93–95^, ECM stiffness-induced alteration in the cell cytoskeleton and stiffness is responsible for many variations in cell signaling and phenotypic changes. Thus, in our model, the parameter (Ecyt), a function of ECM stiffness that reflects cellular stiffness, controls LaminA activation. Figures S1C and D compare *in silico* and experimental variations in cell stiffness^96^ and LaminA activity^53^ for various ECM stiffnesses, respectively. Finally, in Fig. S1E, we compared the model results with six different experimental datasets^55,97–101^ illustrating ECM stiffness-induced variations in YAP/TAZ N/C (nuclear to cytosolic ratio) as a measure of YAP/TAZ activity. These extensive calibration curves establish confidence in our choice of kinetic parameters and model formulation for both chemical and mechanical inputs.

### 2.5 Parametric sensitivity analysis

Next, we determine the sensitivity of the model to different parameters. We chose to conduct a global sensitivity analysis because a local sensitivity analysis has limitations in capturing nonlinear interactions between species in a complex network such as Ca^2+^-mediated YAP/TAZ signaling. Therefore, we performed a Morris global sensitivity analysis^102^ by changing all model parameters (+/- 50% around nominal value), including kinetic and geometric parameters, as well as the initial conditions to compute their global effects on the YAP/TAZ N/C ratio after Tg and Ang II stimulation. We calculated the Morris index (μ*) and standard deviation for each parameter in the model. In Fig 3, we show the model sensitivity analysis results for Tg (panel A) and Ang II (panel B) contexts involving the parameters with high sensitivity scores (μ*>0.05). According to the results, parameters controlling Ca^2+^ channels and pumps, especially SERCA and PMCA, have the highest impact on cell response, followed by geometric and IP_3_-Ca^2+^ module parameters. Given the central role of Ca^2+^ dynamics in both Ang II and Tg contexts, most parameters have similar impacts on cell response in these contexts. However, the parameters that regulate cPKC and PP2A are more specific to Tg and Ang II contexts, respectively.

**Figure 3.**
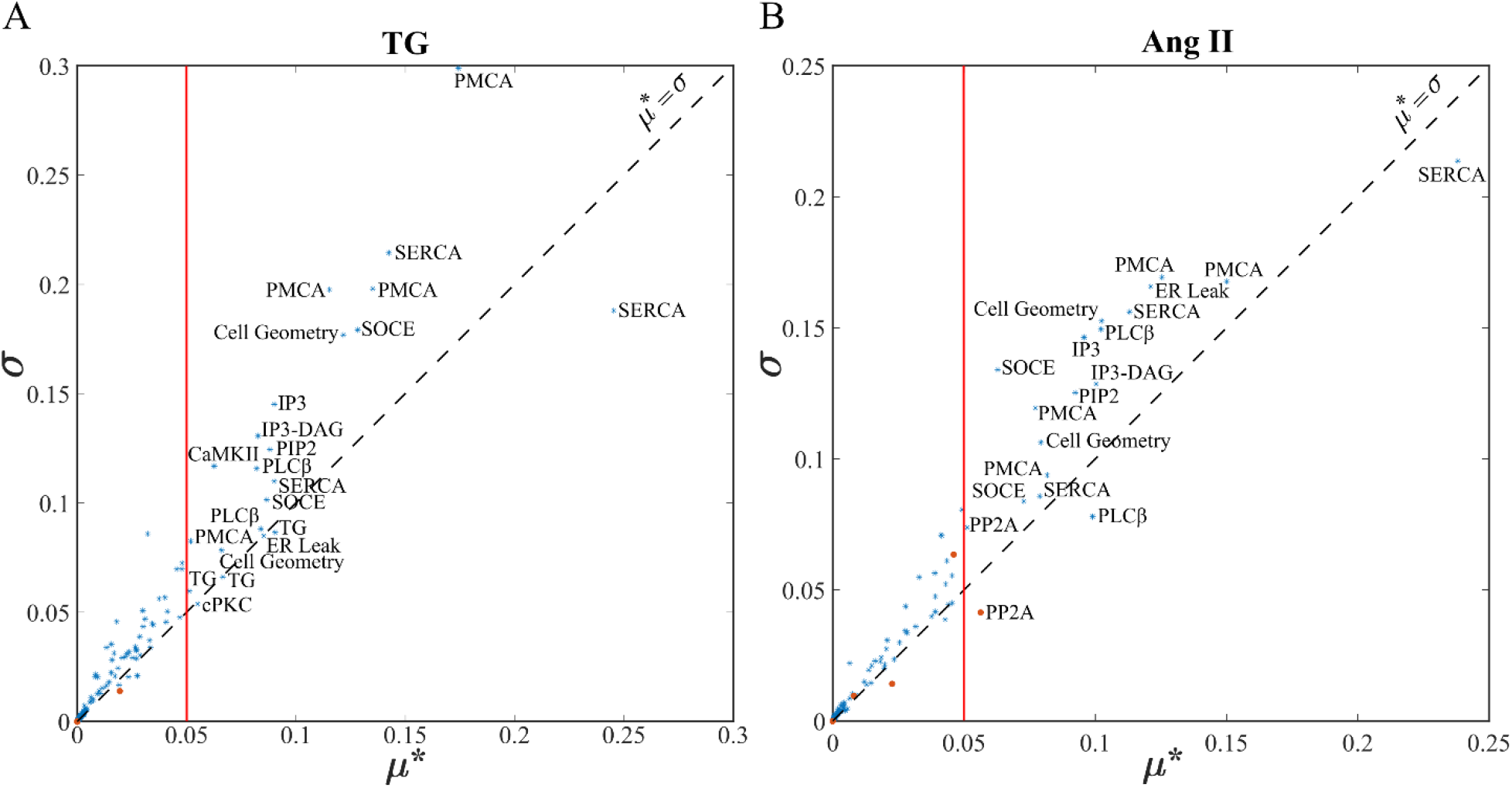
Morris global sensitivity analysis reveals the main parameters of the model regulating Ca^2+^-induced YAP/TAZ response after cell stimulation by 1 μM thapsigargin (Tg) (A) and 0.1 μM Ang II (B). Parameters with μ* > 0.05 are considered influential for Ca^2+^-induced YAP/TAZ activity.

## 3 Results

According to previous experimental observations^27^, there are distinct responses of YAP/TAZ to Ca^2+^ in different experimental settings. This dissimilarity in Ca^2+^-mediated YAP/TAZ response may originate from inherent differences in dynamics and context-dependency of the complex signaling network regulating the YAP/TAZ (Fig. 1A). In this study, by employing the compartmental ODE model of the Ca^2+^-mediated YAP/TAZ signaling, we investigate how the dynamics of Ca^2+^ and signaling modules in the Ca^2+^-mediated YAP/TAZ network and changing the cell context including cell stimuli (type and frequency), and cell type (properties of Ca^2+^ handling proteins) could lead to diverse responses observed for Ca^2+^-YAP/TAZ relationship.

### 3.1 Dynamics of Ca^2+^ and Hippo-YAP/TAZ pathway in response to Ang II and Tg stimulation

We first investigate the dynamics of the Hippo pathway and YAP/TAZ mediated by Ca^2+^. Our input was Tg (1 μM) or Ang II (0.1μM), and stiffness was maintained at zero kPa. Changes in cytosolic Ca^2+^ after Ang II and Tg stimulations are illustrated in Fig. 4A. While Ang II induces a fast Ca^2+^ transient with no increase in steady-state Ca^2+^ level, Tg stimulation leads to a slow Ca^2+^ transient with an elevated level of Ca^2+^ at steady-state. Response of the Hippo pathway, including its core components: MST1/2 and LATS1/2, to changes in cytosolic Ca^2+^ after Ang II and Tg stimulations is displayed in Fig. 4B. MST1/2 is largely dephosphorylated in resting conditions exhibiting only 2-3% of its maximal activity^103^. While no rich time course data is available for MST1/2 after Tg or Ang II stimulation, experimental observations indicate a significant increase in MST1/2 phosphorylation after 30 min stimulation with Tg^28^ and sustained activity of MST1/2 in the presence of Ca^2+ 104^, possibly due to its autophosphorylation^103,105^. Simulations from the model indicate an increase in MST1/2-p from a 2% resting level, a peak between 30 min and 1 hour. and a slow decline (2-5 hours) to a steady-state level higher than its initial level. The model indicates a higher peak value and decline rate for MST1/2 activity after the Tg stimulus (red curve) compared to Ang II (blue curve).

**Figure 4.**
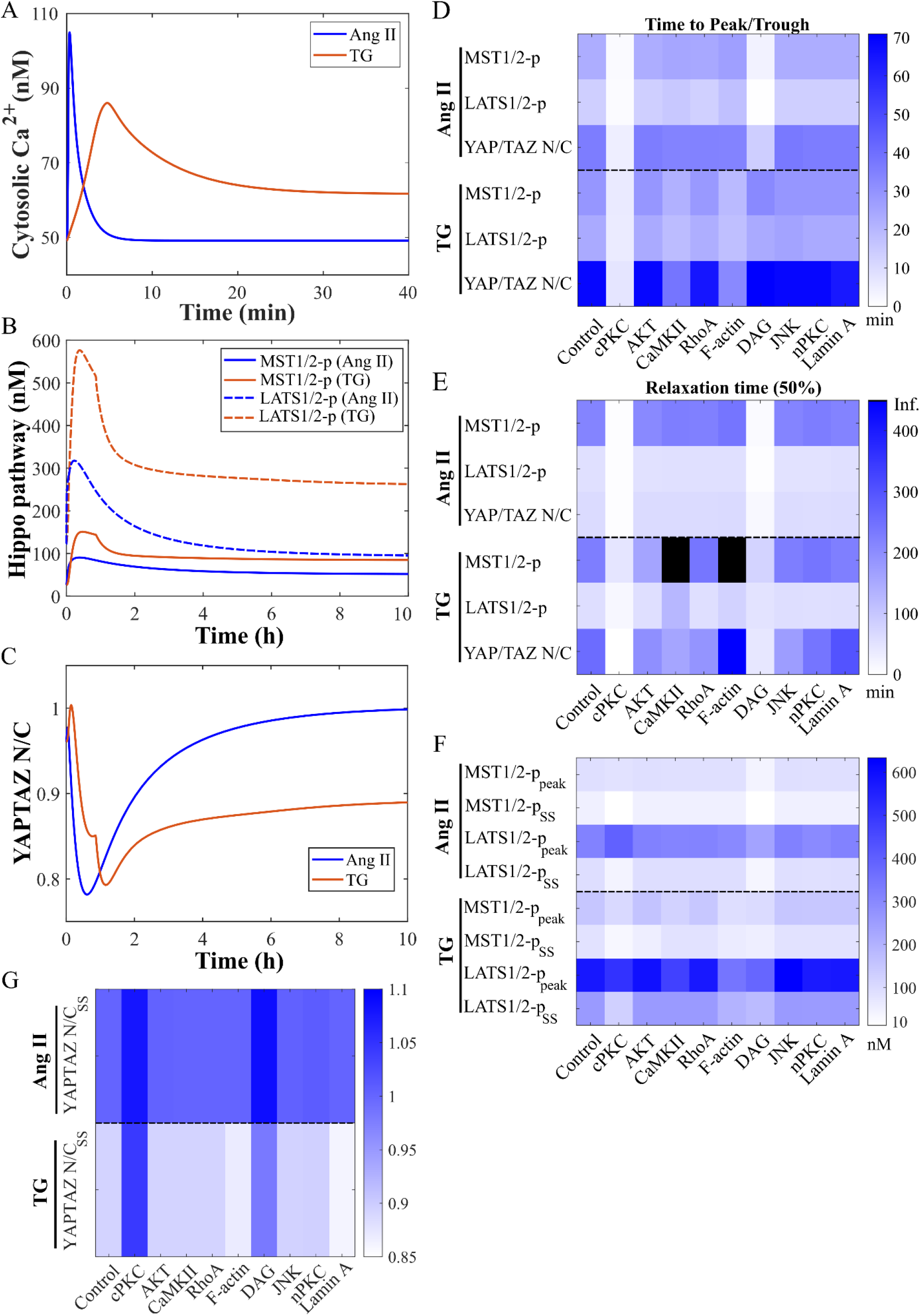
The major mediators in Ca^2+^-mediated YAP/TAZ activation. The dynamic response of Ca^2+^ (A) Hippo pathway core components (B) and YAP/TAZ activity (C) to Ang II and thapsigargin (Tg) stimuli are simulated by the model. In (D-G), the impact of knocking down each upstream regulator on temporal dynamics of Hippo-YAP/TAZ activation (D, E), steady-state and peak values of the Hippo pathway response (F), and steady-state YAP/TAZ activity (G) is shown. The impact of each upstream regulator’s removal is simulated by fixing the regulator’s activity to its initial condition.

We also found that activation trends for LATS1/2 and MST1/2 are similar for both stimuli (Fig. 4B). However, the LATS1/2-p production rate is much larger than MST1/2-p due to multiple upstream activators. Interestingly, in contrast to MST1/2-p, LATS1/2-p has only a higher steady-state level after Tg stimuli when compared to the initial concentration. In the case of Ang II stimulation, the LATS1/2-p steady-state level is lower than its initial level, indicating that Ang II has an inhibitory effect on LATS1/2 in the long term, similar to what has been observed in experimental studies^11,106^.

In the context of Tg stimulus, experimental data indicate an increase in LATS1/2 activity with a peak at 30 min and a fast decline to the initial level^76^. However, the model predicts a higher LATS1/2-p steady-state level than experimental data^76^, partly originating from a higher steady-state level of cytosolic Ca^2+^ after Tg in the simulations. While some studies indicate a higher Ca^2+^ level after Tg^45^, others showed no difference between the initial and steady-state levels of cytosolic Ca^2+^ after Tg stimulation, which could explain low long-term LATS1/2 activity in some experiments^76^.

Next, we study the impact of Ang II and Tg on YAP/TAZ activity, reported as YAP/TAZ N/C ratio^36,50^. As shown in Fig. 4C, after a small and fast uprise, YAP/TAZ N/C ratio decreases to its minimum (∼30-40 min for Ang II and 60-70 min for Tg) and then rises to reach its steady-state value after 10 hours. Interestingly, while both Ang II and Tg result in elevated Ca^2+^ levels, their YAP/TAZ activities in the long term are significantly different, indicating the important role of cell stimuli (context) in diverse Ca^2+^-YAP/TAZ relationship^27^.

### 3.2 Context-dependent contribution of upstream regulators to Hippo-YAP/TAZ activity

We next explored the role of upstream regulators from different signaling modules, including cPKC, AKT, CaMKII, RhoA, F-actin, DAG, JNK, nPKC, and LaminA, on the dynamics of the Hippo pathway and YAP/TAZ. To do this, we simulated the removal of each regulator’s effect on its downstream targets by setting the regulator’s rate of variations equal to zero. Then, we simulated the variations of MST1/2-p, LATS1/2-p, and YAP/TAZ N/C levels in both Ang II and Tg contexts. This analysis allowed us to determine the degree that each upstream pathway impacts the Hippo pathway and YAP/TAZ activity. We determined the impact of removing each regulator on time to peak or trough (Fig. 4D) and relaxation time (Fig. 4E) for MST1/2-p, LATS1/2-p, and YAP/TAZ activity after Tg and Ang II stimulation. Removal of cPKC or DAG makes the dynamics of Hippo and YAP/TAZ faster by decreasing peak/trough and relaxation times. We then plotted the steady-state concentrations and peak values of MST1/2-p and LATS1/2-p after removing each upstream regulator effect for both Ang II and Tg stimuli in Fig. 4F. We found that removing the contribution of cPKC decreases active MST1/2 and LATS1/2 steady-state levels as well as active MST1/2 peak levels in both contexts. However, cPKC or DAG removal increases the active LATS1/2 peak level. In addition, by comparing steady-state levels of YAP/TAZ activity after removing the contribution of each regulator (Fig. 4G), we found that PKC removal also results in higher steady-state YAPTAZ activity. In brief, Ca^2+^-increase in cPKC or DAG activity could make the Hippo and YAP/TAZ dynamics slower and leads to lower YAP/TAZ activity in the long term through elevating MST1/2 and LATS1/2 steady-state phosphorylation.

CaMKII is a known Ca^2+^ downstream target affecting many signaling pathways^66,107,108^. Our model predicts that removing the effect of CaMKII reduces MST1/2-p and LATS1/2-p peak levels only after Tg, not Ang II (Fig. 4F). This stimulus-specific response is more apparent in its effect on Hippo pathway dynamics. While abolishing the CaMKII contribution reduces the peak times of the Hippo pathway in the Tg context, it slightly increases these times in the Ang II context (Fig.4D-E). It also significantly increases Hippo pathway relaxation times for Tg stimulus without any significant impact on relaxation times for Ang II stimulus. Regarding YAP/TAZ, CaMKII contribution’s removal only significantly decreases the trough and relaxation times for the Tg stimulus and has no effects on YAP/TAZ steady-state levels for both stimuli (Fig.4D-G). Thus, the model predicts that CaMKII contribution to Hippo and YAP/TAZ activity is more pronounced in cell stimulation by Tg rather than Ang II. Moreover, the CaMKII impact is more limited to Hippo and YAP/TAZ temporal dynamics than steady-state activities.

F-actin is a key cytoskeletal component implicated in the regulation of YAP/TAZ nuclear translocation^55,109^. Removal of F-actin contribution to Ca^2+^-mediated Hippo-YAP/TAZ pathway activity results in the decrease in Tg-induced MST1/2-p and LATS1/2-p peak and steady-state levels and Ang II-induced LATS1/2-p peak level (Fig.4F). It also diminishes peak/trough times of the Hippo-YAP/TAZ pathway in the Tg context and increases peak times of the Hippo pathway in the Ang II context (Fig. 4D). The removal of F-actin contribution increases relaxation times of MST1/2-p and YAP/TAZ only in Tg context. Besides, it only decreases Tg-induced YAP/TAZ steady-state level (Fig. 4G). In summary, F-actin can participate in both the inhibition and activation of YAP/TAZ. On one hand, higher F-actin increases YAP/TAZ nuclear translocation through cytosolic stiffness modulation. On the other hand, it can contribute to YAP/TAZ inhibition by cPKC-dependent activation of LATS1/2. Given these two opposite effects, the model predicts that F-actin in the long-term activates YAP/TAZ after a Tg-induced increase in Ca^2+^.

LaminA controls YAP/TAZ nuclear translocation by inducing the stretching of NPCs resulting in their lower resistance against YAP/TAZ nuclear import. As expected, removal of LaminA contribution only affects YAP/TAZ dynamics but not MTS1/2 or LATS1/2 dynamics. Lack of LaminA changes in our model decreases the time to trough for YAP/TAZ and the YAP/TAZ steady-state level only after Tg stimulus and elevates Tg-induced YAP/TAZ relaxation time. While other regulators like AKT, RhoA, JNK, and nPKC contribute to the dynamics of the Hippo-YAP/TAZ pathway at some level (Figs. 4D-F), their effects are significantly lower than regulators discussed above. In summary, we find that while the dynamics of YAP/TAZ falling to its trough is mainly controlled by phosphorylation, nuclear translocation regulates its rise to the steady-state level.

### 3.3 Ca^2+^ temporal dynamics govern the switching of the Ca^2+^-YAP/TAZ relationship

One of the potential causes of diversity in the Ca^2+^-YAP/TAZ relationship is the significant variation of the cytosolic Ca^2+^ dynamics observed in different cell contexts (e.g., cell types and experimental settings)^27,39,110^. These variations are occasionally unrecognizable due to the noise or Ca^2+^ waves but may result in distinct downstream responses like Ca^2+^-YAP/TAZ diverse relationships. In this section, we explore how different transient Ca^2+^ dynamics may lead to Ca^2+^-induced activation or inhibition of YAP/TAZ. We generated several artificial Ca^2+^ transients by perturbing two model parameters with the highest impact on Ca^2+^-mediated YAP/TAZ response identified from Morris sensitivity analysis (see Fig.3). These parameters are Michaelis constant (Km) for PMCA function and Hill power (n) for SERCA function. Figures 5A-I, III illustrate the Ca^2+^ transient response in the Tg context by altering n (SERCA) and Km (PMCA) around their nominal values of 1 and 0.2 μM, respectively. While altering n (SERCA) significantly modifies Ca^2+^ initial resting level and peak time, its effect on Ca^2+^ steady-state level, relaxation time, and the peak value is negligible (Fig. 5A I). Therefore, we observed significant changes in initial YAP/TAZ activity and its dynamics in the first hours after stimulation (Fig. 5A-II). However, its steady-state level and dynamics at longer times are nearly unchanged. This may have an impact on the direction of changes in YAP/TAZ activity after the elevation of cytosolic Ca^2+^ in the cell. According to model results, a Ca^2+^ transient with and without steady-state elevation in Ca^2+^ concentration could lead to the opposite direction of changes, increase or decrease, in YAP/TAZ steady-state activity. Thus, the ratio of steady-state to the initial level of cytosolic Ca^2+^ during Ca^2+^ stimulation could predict the Ca^2+^-YAP/TAZ relationship in the cell.

**Figure 5.**
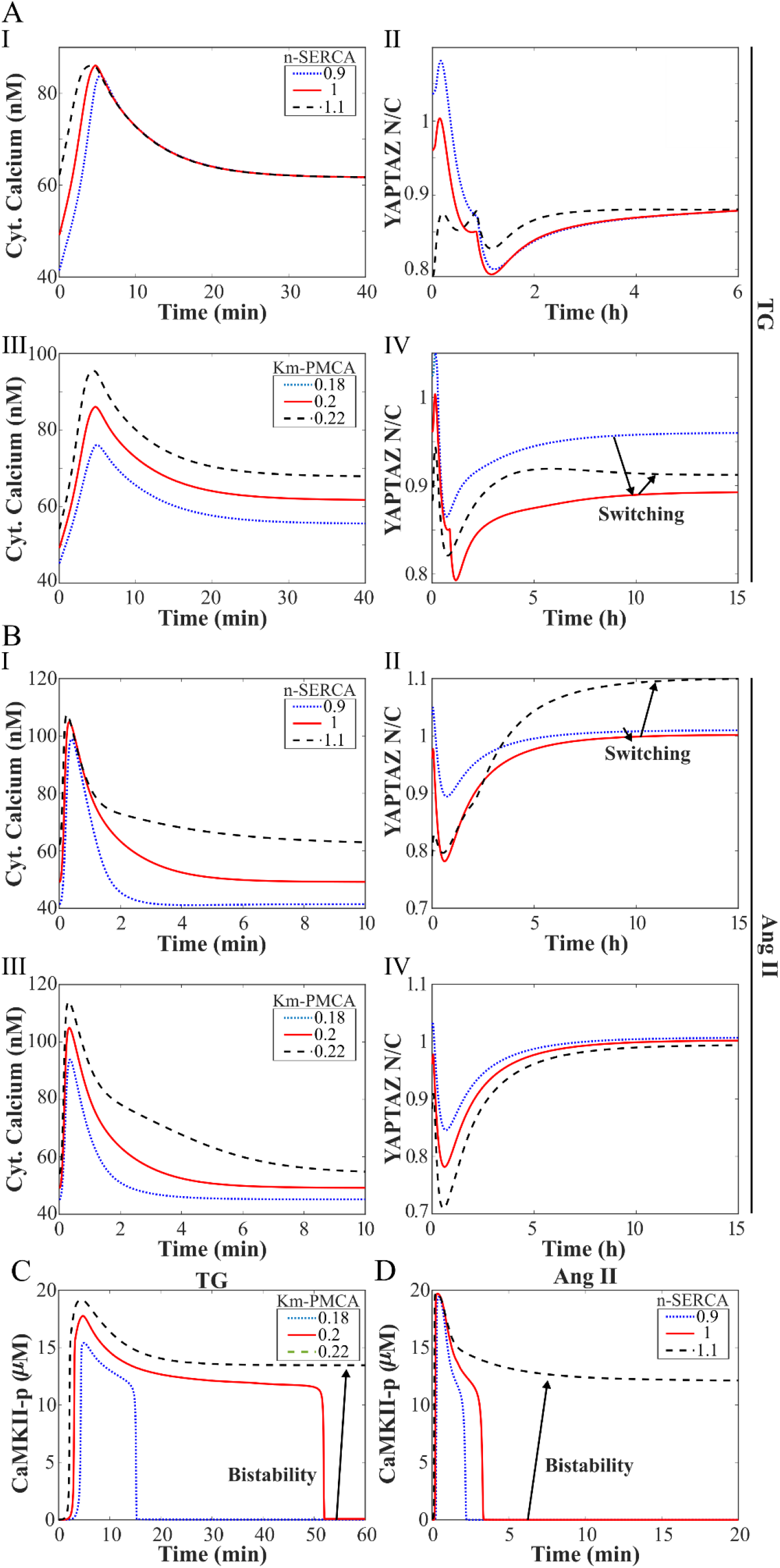
Impact of Ca^2+^ dynamics on YAP/TAZ activity. (A) Time course of Ca^2+^ transient and following YAP/TAZ activity for different n-SERCA (I, II) and Km-PMCA (III, IV) values in the context of Tg stimulus. (B) Time course of Ca^2+^ transient and following YAP/TAZ activity for different n-SERCA (I, II) and Km-PMCA (III, IV) values in the context of Ang II. (C) Bistable response of CaMKII after varying Km-PMCA in the Tg context. (D) Bistable response of CaMKII after altering n-SERCA in the Ang II context.

Varying Km-PMCA has a significantly different impact on the Ca^2+^ transient in comparison with altering n-SERCA. Change in Km-PMCA shifts the entire Ca^2+^ transient response to higher or lower Ca^2+^ levels without any change in peak and relaxation times or any significant changes in the ratio of steady-state to the basal level of cytosolic Ca^2+^ (Fig. 5A-III). The resultant YAP/TAZ activity tends to follow this shift for a longer time scale. However, as the Ca^2+^ time course shifts toward higher Ca^2+^ levels, the Ca^2+^-YAP/TAZ relationship switches from inhibition to activation. This result predicts that the absolute level of cytosolic Ca^2+^ could also govern YAP/TAZ bidirectional response.

Another switching response was observed after modifying the n-SERCA parameter after Ang II stimulation. As shown in Fig. 5B-I, increasing n-SERCA in the Ang II context results in a significant increase in Ca^2+^ initial and steady-state levels as well as higher relaxation time without significant change in peak value.

Interestingly, a switching event reversed this trend instead of seeing a shift to lower YAP/TAZ transients because of higher Ca^2+^ levels, as observed in Fig. 5B-IV after varying Km-PMCA in the Ang II context. However, if we compare the ratio of steady-state to initial YAP/TAZ, we observe a smooth change in YAP/TAZ activity from inhibition to activation. It is because of a higher decrease in YAP/TAZ initial level than the steady-state level after increasing Km-PMCA in Fig. 5B-IV. These results confirm the previous model predictions on the contribution of higher Ca^2+^ levels to the change in the Ca^2+^-YAP/TAZ relationship. However, the difference between YAP/TAZ responses after the Ang II stimulus (Fig. 5B) despite similar Ca^2+^ transient predicts the high sensitivity of YAP/TAZ temporal response to Ca^2+^ transient characteristics such as Ca^2+^ amplitude. In summary, model results predict the significant impact of cytosolic Ca^2+^ levels at the initial and steady-state (relative and absolute) on the Ca^2+^-YAP/TAZ relationship and the impact of Ca^2+^ amplitude on YAP/TAZ temporal response.

To explore the cause of the switching phenomenon that may explain different Ca^2+^-YAP/TAZ relationships, we screened the variations of all signaling components in the model and found the CaMKII bistable response as a potential source of this switching event. As shown in Figures 5C and D, varying Km-PMCA in the Tg context and n-SERCA in the Ang II context led to a change from low to high steady-state CaMKII activity followed by YAP/TAZ switching. These results predict that if we have a bistable response of CaMKII as observed in some cell types^111,112^, high CaMKII activity could lead to a positive Ca^2+^-YAP/TAZ relationship.

### 3.4 YAP/TAZ exhibits a non-monotonic response to periodic GPCR activation

Repetitive transient increases in IP_3_ and cytosolic Ca^2+^ after GPCR stimulation can be seen in a broad range of non-excitable cells^113^. However, it is not clear how these periodic activations of the IP_3_-Ca^2+^ module affect the YAP/TAZ pathway. To explore the sensitivity of YAP/TAZ activity to the frequency of IP_3_-Ca^2+^ module activation, we simulated Ang II-induced activation of the IP_3_-Ca^2+^ module using a square wave input (Ang II) at various periods (T) from 2 days to 10 min. Each period involves two equal times for cell activation and rest. For example, in a 2 days period, we have one day of Ang II stimulation and one day of cell rest. In the 2 days period, there is sufficient time for YAP/TAZ signaling components to reach their steady-state level without any frequency-related impact on YAP/TAZ (Fig. 6A). However, by lowering the period (T) from 2 days to 10 min, we observed a steady decrease in YAP/TAZ amplitude (difference between the mean value and max/min in each period) and a non-monotonic response for YAP/TAZ mean activity (Figs. 6B-C). Lowering the period (T) from 2 days to 2 h elevated the YAP/TAZ mean activity. But a further decrease from 2 h to 10 min reduced YAP/TAZ mean activity resulting in a maximum for YAP/TAZ mean activity in the 2 h period. By comparing steady-state levels of YAP/TAZ between 2 days and 10 min, we can predict that increasing the frequency could change the direction of the Ca^2+^-YAP/TAZ relationship in the Ang II context from a positive relationship to a negative one.

To investigate which signaling mediators participate in this frequency-dependent YAP/TAZ response, we simulated the response of main mediators in the model to periodic GPCR activation by Ang II for 2 days, 2 h, and 10 min periods. All mediators showing significant frequency-dependent effects are illustrated in Figs.6D-I. The JNK-p and AKT-p inhibit the Hippo pathway and can thus be considered activators of YAP/TAZ. As shown in Figures 6D-E, a decrease in period (T) reduces the amplitude of both JNK-p and AKT-p. However, JNK-p and AKT-p responses are completely different regarding the direction of changes in the mean activity and period threshold. Based on the results, we estimated a period threshold between 2 days and 2h for JNK-p and a lower period threshold between 2h and 10 min for AKT-p. Pyk2-p activates the RhoA pathway and then F-actin; it can be considered a YAP/TAZ activator (Fig. 1C). A decrease in period (T) results in a lower amplitude and higher mean value for Pyk2-p (Fig. 6F).

**Figure 6.**
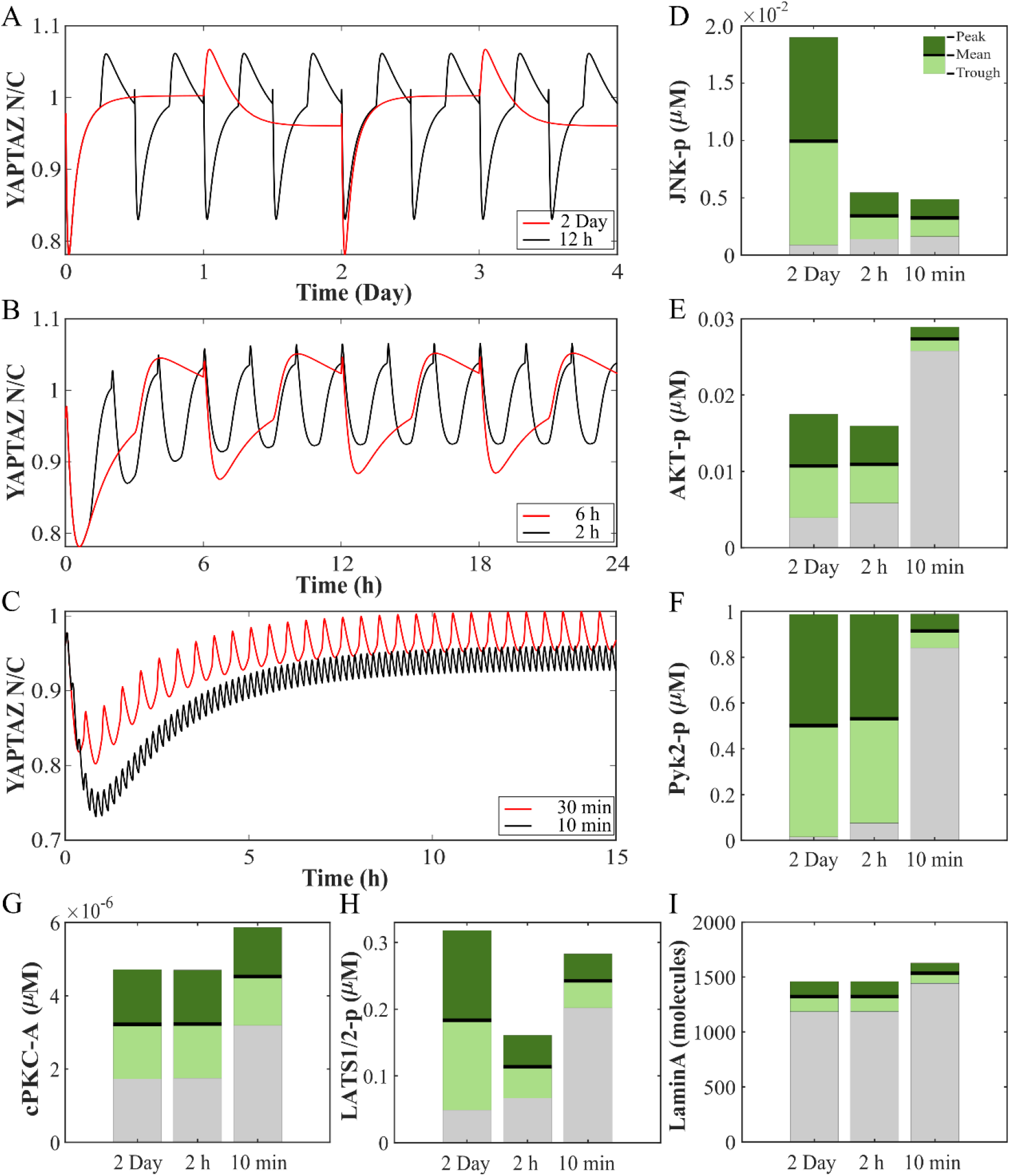
The YAP/TAZ exhibits a non-monotonic response to periodic GPCR activation. (A-C) YAP/TAZ activation time course for stimulation periods from 2 Days to 10 min. The oscillation domain (red and blue bars) and the mean activity (thick black line) of upstream regulators of YAP/TAZ, including JNK (D), AKT (E), Pyk2p (F), cPKC (G), LATS1/2 (H), and Lamin A (I) are shown for stimulation periods from 2 Day to 10 min.

cPKC has a bidirectional effect on the Hippo pathway^33,114^. However, our results (Fig. 4F) indicated a dominant cPKC inhibitory action on YAP/TAZ steady-state activity. As shown in Fig. 6G, a decrease in the period T elevates the mean activity of cPKC without significantly changing its amplitude. LATS1/2, as a direct inhibitor of YAP/TAZ, significantly contributes to its frequency-dependent response. As shown in Fig. 6H, the LATS1/2-p response is the opposite of the YAP/TAZ response in terms of mean activity. Finally, LaminA, an activator of YAP/TAZ that facilitates its nuclear translocation, exhibits a slight increase in the mean activity after a decrease in period T.

In summary, model results predict a non-monotonic YAP/TAZ activity in terms of period T due to the distinct response of YAP/TAZ upstream regulators to periodic GPCR activation. The model predicts that while frequency-induced inhibition of JNK and activation of cPKC favor a decrease in YAP/TAZ activity, activation of AKT, Pyk2, and LaminA tend to increase YAP/TAZ activity. Accordingly, the results predict the dominance of cPKC and JNK impact on YAP/TAZ in longer periods and AKT, Pyk2, and Lamin A impacts in shorter periods. Results also confirmed a significant role for LATS1/2 in mediating YAP/TAZ frequency-dependent response.

## 4 Discussion

Here, we developed a prior knowledge-based network model of Ca^2+^-mediated YAP/TAZ signaling to examine how different temporal dynamics of signaling species contribute to distinct YAP/TAZ responses observed in experiments. The model comprises seven signaling modules and their crosstalk that interact with Ca^2+^ and transduce biochemical (Ang II) and biomechanical signals (ECM stiffness) to YAP/TAZ. The identified signaling modules include GPCR, IP_3_-Ca^2+^, Kinases, RhoA, F-actin, Hippo, and YAP/TAZ module. The model captures both time course and steady-state data available in the literature. By employing our model, we make a series of experimentally testable predictions. First, we predicted that while both Ang II and Tg stimuli decrease YAP/TAZ activity in the short time, AngII activates YAP/TAZ in the long term in contrast to Tg. Second, we identified the major mediators of Ca^2+^-induced YAP/TAZ activation (e.g., cPKC, DAF, CaMKII, F-actin, and LaminA) and predicted their context-specific contribution to YAP/TAZ dynamics and steady-state behavior. Third, we predicted how variations in Ca^2+^ transients in different cell contexts might lead to the controversial Ca^2+^-YAP/TAZ relationship observed in experimental data. We predicted the relationship between basal and steady-state cytosolic Ca^2+^ levels and direction of changes in Ca^2+^-induced YAP/TAZ activity, the contribution of Ca^2+^ amplitude on YAP/TAZ temporal response, and the significant role of CaMKII bistability in switching Ca^2+^-YAP/TAZ relationship. Finally, we predicted a non-monotonic YAP/TAZ activity in response to periodic GPCR activation because of distinct frequency-dependent responses of YAP/TAZ upstream regulators.

### 4.1 The cPKC and Hippo core kinases integrate signals with distinct temporal scales in YAP/TAZ regulation

Cells experience and respond to numerous biochemical and mechanical stimuli during their lifetime. Although multiple mechanisms have been identified for cell mechanotransduction, a major part of these mechanisms act through ECM and lead to cell cytoskeletal remodeling in the order of several hours to days^115^. Unlike mechanical activators like ECM stiffness, a majority of biochemical stimuli like GPCR agonists are considered fast inputs resulting in receptor activation and downstream signaling in the order of seconds to minutes^11,116^. For proteins like YAP/TAZ sensitive to both fast and slow stimuli^6,16^, upstream regulators that integrate signals with different time scales are necessary. In Ca^2+^-mediated YAP/TAZ signaling, we predict that cPKC and core Hippo pathway components (MST1/2 and LATS1/2) may play this role. Removing the effects of cPKC activation in Ang II and Tg contexts significantly shifts the Hippo-YAP/TAZ dynamics to a faster response with a lower transient peak for Hippo pathway components and higher steady-state activation of YAP/TAZ.

Inhibition of cPKC as one of the key effectors downstream of GPCR can activate or inhibit YAP/TAZ in a context-dependent manner^27,33,73^. cPKC inhibition effectively blocked dephosphorylation of YAP/TAZ by acetylcholine (Gq11-coupled receptor agonist) in U251MG cells^33^. It also blocked amlodipine- or ionomycin-induced YAP/TAZ phosphorylation in LN229 cells^28^. In addition to cPKC, the sustained activity of the Hippo pathway, especially MST1/2, after Ang II significantly contributes to the YAP/TAZ sensitivity to fast GPCR signals. Furthermore, AKT has the potential for integrating upstream signals, as shown in Fig. 6. However, AKT removal did not noticeably affect MST1/2 and LATS1/2 peak and steady-state levels based on model results. Based on our predictions, we can hypothesize that while MST1/2 phosphorylation by AKT can decrease MST1/2 activity^67,103^, because of the large unphosphorylated portion of MST1/2 in cells^103^, Ca^2+^-induced AKT activation cannot practically limit activation of MST1/2 by cPKC and F-actin.

### 4.2 CaMKII autonomous activity controls Ca^2+^-YAP/TAZ bidirectional relationship

CaMKII is an auto-phosphorylating kinase that acts as major Ca^2+^ signaling downstream in cell physiology and pathology^112^. Autophosphorylation of CaMKII could lead to Ca^2+^/CaM-independent (autonomous) kinase activity that can last 30 min or longer after a short-term transient increase in cytosolic Ca^2+^ concentration^117^. To cover Ca^2+^ signaling in various cell types, our model considers both Ca^2+^/CaM-dependent and autonomous activation of CaMKII (Table 3). As shown in Fig. 5, a change in the Ca^2+^ dynamics could alter the direction of changes in Ca^2+^-YAP/TAZ interaction from Ca^2+^-induced inhibition of YAP/TAZ to activation, and CaMKII autonomous activation could mediate this switching process.

In the context of Tg stimulation, the model predicts a high activity for CaMKII starting from ∼5 min and ending before one hour. Likewise, Timmins et al.^118^ illustrated that after Tg stimulation, CaMKII rises to its maximum in 5 min and declines to its initial level in 30 min. Zhong et al.^85^ also reported Tg-induced CaMKII activation with a steady increase till 30 min and then return to the resting level before one hour. For the Ang II context, the model predicts a rise to maximum CaMKII activity in 1 min and a return to the initial level in less than 5 min. Zhu et al.^119^ showed similar activation of CaMKII after Endothelin-1, a Gq activator, with maximum CaMKII activity in 1 min and returning to the initial level in 6 min. However, the Ang II-induced CaMKII transient response can be switched to a sustained response (Fig. 5D) with high CaMKII activity even after one day^120^. Given the significant variability between cell types in Ca^2+^ dynamics and CaMKII autonomous activity, the model results predict that in the cells with potential CaMKII autonomous activity, the context-dependent duration of high CaMKII activity could alter the Ca^2+^-YAP/TAZ relationship from one context to another^27^. In the cells where CaMKII autonomous activity is not significant, we expect a more robust cell response in terms of the Ca^2+^-YAP/TAZ relationship to variations in Ca^2+^ transient, especially for changes reflecting a shift of Ca^2+^ transient to higher or lower concentrations (Fig. 5A-III).

### 4.3 The model predicts Ca^2+^-YAP/TAZ distinct relationships in experiments

The Ca^2+^-YAP/TAZ relationship is controversial. Multiple experimental studies reported both inhibition and activation relationship between Ca^2+^ and YAP/TAZ reviewed by Wei et al.^27^. To explore the cell response and Ca^2+^-YAP/TAZ relationship in each study, we simulated various experimental conditions by the model. The model results are compared with experimental data in Table 5. In the first study, authors knocked out the TPCN2 gene in CHL1 and B16-F0 (murine primary melanoma) cell lines and observed activation of YAP/TAZ in the cells^31^. They suggested the contribution of Orai 1 and PKCβ in the activation of YAP/TAZ, given the significant decrease in their expressions in the knockout cells. We simulated this experiment with a 30% decrease in Ymax_SO_ value (see Table 2, J_SOCE_) and a 90% decrease in cPKC initial concentration (Table 1) to reproduce reported experimental changes in Orai1 and cPKC expression^31^. As shown in Table 5, the model predicts an increase in YAP/TAZ activity similar to experimental data. Interestingly, while simulating each perturbation at a time increased the YAP/TAZ N/C ratio, applying both perturbations together yielded less of an increase in YAP/TAZ N/C than just cPKC knockdown, indicating the non-linearity of cell response in Ca^2+^-mediated YAP/TAZ activity.

**Table 5.**
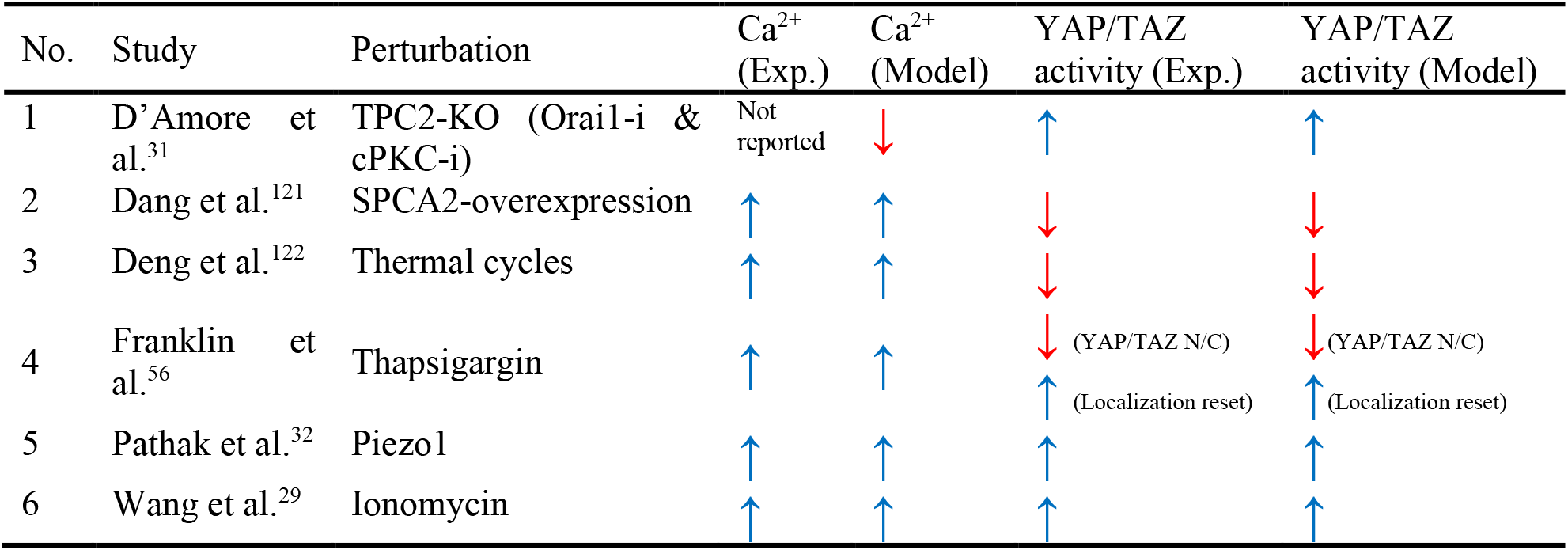
Prediction of Ca^2+^-YAP/TAZ relationship in different experimental settings

In the second study, Dang et al.^121^ showed that overexpression of Ca^2+^-ATPase isoform 2 (SPAC2) in breast cancer cells elevates baseline Ca^2+^ and YAP phosphorylation and decreases YAP/TAZ activity. Consistent with the experiment, the model predicts an increase in YAP phosphorylation and a decrease in YAP/TAZ activity after a 30% increase in baseline Ca^2+^ by applying higher extracellular Ca^2+^. The third study demonstrates that upregulation of Ca^2+^ influx and higher cytosolic Ca^2+^ levels in hADSC cells after thermal cycles decrease YAP nuclear localization (activity)^122^. Given the role of TRP channels in the thermal-induced changes in Ca^2+^ influx^122^, we simulated the experiment by cyclic activation of the TRP channel in the model with a similar temperature cycle reported for hADSC cells^122^. The model predicts a cyclic response for YAP/TAZ with a range of activities lower than its initial.

In a study by Franklin et al.^56^, authors showed a rapid YAP localization reset after Tg stimulation, including an initial fast depletion of nuclear YAP followed by slow nuclear enrichment on the time scale of 0.5–2h with a steady-state YAP/TAZ N/C mean value lower than its initial. As shown in Fig. 4A, the model predicts similar YAP localization reset in around 2 hours leading to a decreased steady-state YAP/TAZ N/C. While steady-state YAP N/C is reduced in both experiment and model, Franklin et al.^56^ indicated higher expression of YAP target genes after Tg stimulation and linked higher gene expression to the reentry of YAP to the nucleus. This observation highlights the significance of YAP/TAZ temporal dynamics in regulating cellular functions. Thus, part of the reported diversity in the Ca^2+^-YAP/TAZ relationship may originate from differences between studies in measuring YAP/TAZ activity based on changes in the steady-state YAP/TAZ N/C or expression of YAP/TAZ target genes.

In another study, Pathak et al.^32^ found that ECM stiffness-induced Ca^2+^ influx by Piezo1 is required for Y AP nuclear localization when human neural stem cells (hNSPCs) grow on a stiff surface. They observed that hNSPCs transfected with Piezo1 siRNA displayed nuclear exclusion more frequently than cells transfected with non-targeting siRNA when grown on glass coverslips^32^. To reproduce the experiment conditions, we simulated YAP/TAZ N/C after ECM stiffness stimulus (glass∼ 70 GPa range) with and without TRP channel knockdown. In the model, the TRP channel is responsible for sensing ECM stiffness-induced changes in cytosolic Ca^2+^, similar to the observed Piezo1 function in hNSPCs. Consistent with the experiment, the model predicts a significant reduction in ECM stiffness-induced YAP/TAZ N/C increase after the knockdown of the TRP channel. Finally, Wang et al.^29^ displayed that Ca^2+^ ionophore ionomycin increases TAZ in human hepatocytes. We simulated this experiment by reproducing iono-induced Ca^2+^ transient^123,124^ via a short-time (30 s) increase in Ca^2+^influx through the plasma membrane (doubling Kf_PM_ parameter; see Table 2), which results in a Ca^2+^ transient similar to Ang II-induced Ca^2+^ (Fig. 4A). The model predicts an increase in RhoA (2h) and TAZ activities (4h and later) after ionomycin agreeing with the experimental observations.

### 4.4 Future directions

In our model, although we calibrated the model to capture ECM stiffness-induced cell response to simulate experiment conditions, fully understanding the impact of concurrent biochemical and biomechanical stimuli on the Ca^2+^-YAP/TAZ relationship requires more time-course data and spatial modeling^125^. Moreover, the varied 3D geometry of cells could also be one of the factors leading to the divergent Ca^2+^-YAP/TAZ relationship. As discussed in Scott et al. study^36^, three-dimensional (3D) environments made the YAP/TAZ activation by stiffness uncertain. Thus, a computational model considering both temporal and spatial dynamics of YAP/TAZ regulations is desired to predict the impacts of concurrent stimuli as well as cell geometry on the Ca^2+^-YAP/TAZ relationship. Accordingly, in a future study, we aim to provide a comprehensive model of YAP/TAZ capturing both temporal and spatial dynamics of Ca^2+^-mediated YAP/TAZ activation.

### Summary

Experimental findings suggest that Ca^2+^ could be an intracellular messenger for the Hippo-YAP/TAZ pathway in response to biomechanical and biochemical stimuli. However, studies reported contradictory Ca2+-YAP/TAZ relationships in different cell contexts, making it difficult to unravel the role of Ca^2+^ in YAP/TAZ-dependent cellular functions from a purely experimental approach. In this study, we developed a network model of Ca^2+^-mediated YAP/TAZ signaling and investigated underlying mechanisms regulating the Ca^2+^-YAP/TAZ relationship, as observed in experiments. The model predicted the role of Ca^2+^ temporal dynamics, CaMKII autonomous activity, and stimulus frequency in modifying the Ca^2+^-YAP/TAZ relationship. The model predicted Ca^2+^-YAP/TAZ distinct relationships in different settings consistent with experiments. This model provides a coherent framework to predict the cell response to biochemical and biomechanical stimuli integrated through Ca^2+^ signaling.

## List of Abbreviations

Abbreviation: Meaning
AKT: protein kinase B
Ang II: Angiotensin II
Arp2/3: Actin-related protein 2/3 complex
CaMKII: Calcium/calmodulin-dependent protein kinase II
DAG: Diacylglycerol
ECM: Extracellular matrix
ER: Endoplasmic reticulum
FAK: focal adhesion kinase
GPCRs: G-protein coupled receptors
IP_3_: Inositol 1,4,5-trisphosphate
IP_3_R: IP_3_ receptor
JNK: c-Jun N-terminal kinase
LATS1/2: Large tumor suppressors 1 and 2
LIMK: LIM kinase
mDia: mammalian diaphanous-related formin
Merlin: Moesin-ezrin-radixin like
MST1/2: Mammalian STE20-like protein kinase 1/2
Myo: myosin light chain
NPCs: nuclear pore complexes
PA: Phosphatidic acid
PIP2: Phosphatidylinositol 4,5-bisphosphate
PKC: protein kinase C
PLCβ: Phospholipase C β
PLD: Phospholipase D
PMCA: Plasma membrane Ca^2+^ ATPase
PP2A: Protein Phosphatase 2A
Pyk2: protein tyrosine kinase 2
RhoA: Rho family of GTPases
ROCK: Rho-associated kinase
RyRs: Ryanodine receptors
SERCA: Sarco/endoplasmic reticulum Ca2+-ATPase
SOCE: Store-operated calcium entry
TAZ: Transcriptional coactivator with PDZ-binding motif
TEADs: TEA domain transcription factors
Tg: Thapsigargin
TPC2: Two-pore channel 2
TRPV4: Transient receptor potential vanilloid-type 4
YAP: Yes-associated protein

## Acknowledgments

This work was supported in part by the Wu Tsai Human Performance Alliance at UCSD to SIF and PR. AK was also supported by an AHA postdoctoral fellowship.

## Supplementary materials

**Figure S1.**
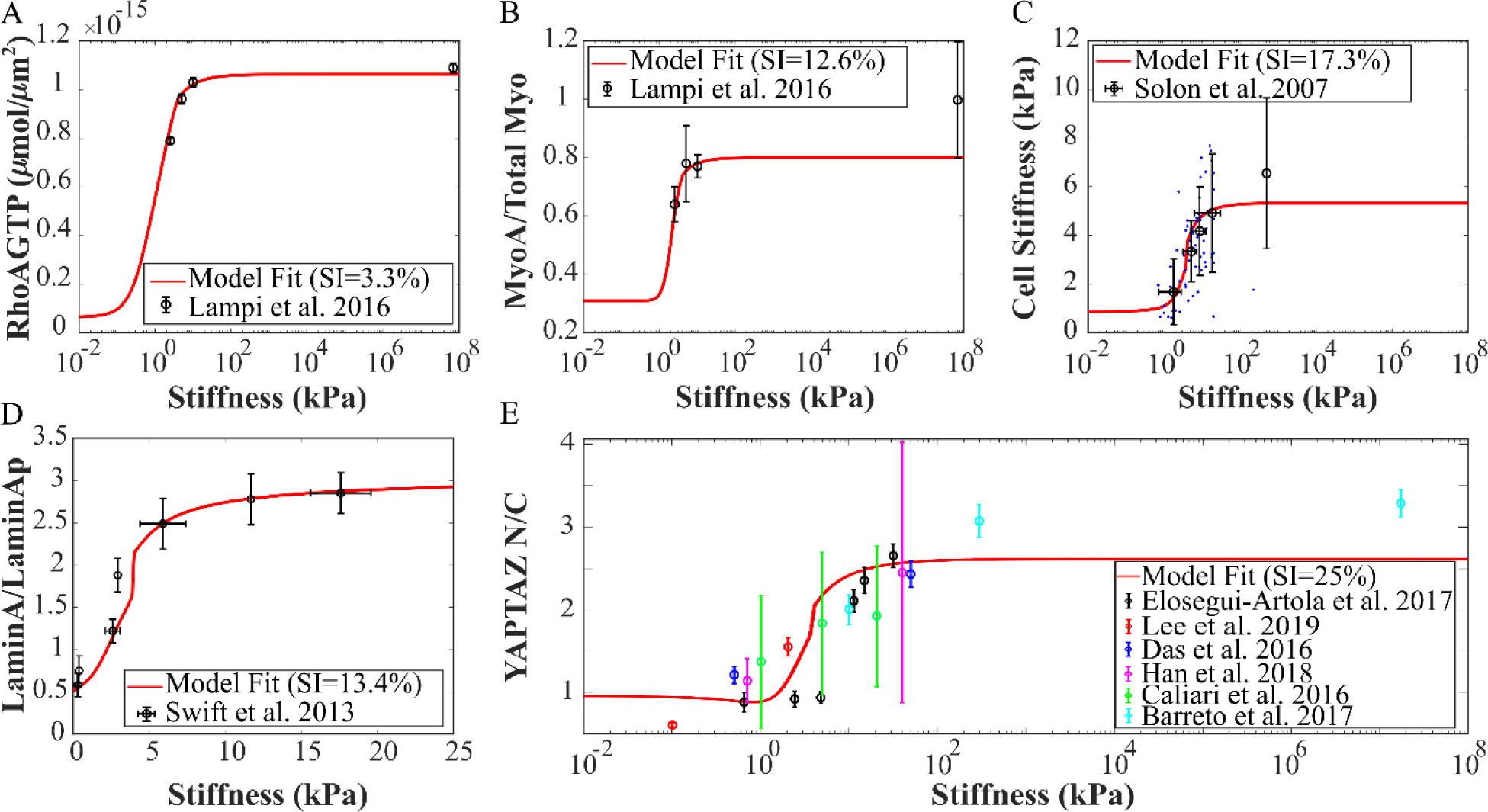
The compartmental ODE model captures the steady-state response of ECM stiffness-induced YAP/TAZ regulation in the cell. Model parameters are estimated to capture steady-state variations of RhoA-GTP (A), myosin (B), cell stiffness (C), Lamin A (D), and the ratio of nuclear YAP/TAZ to cytosolic YAP/TAZ (E) for a range of substrate stiffnesses.

