## Supplementary figures and images for "Mechanisms underlying divergent relationships between Ca^2+^ and YAP/TAZ signaling"

### Fig S1

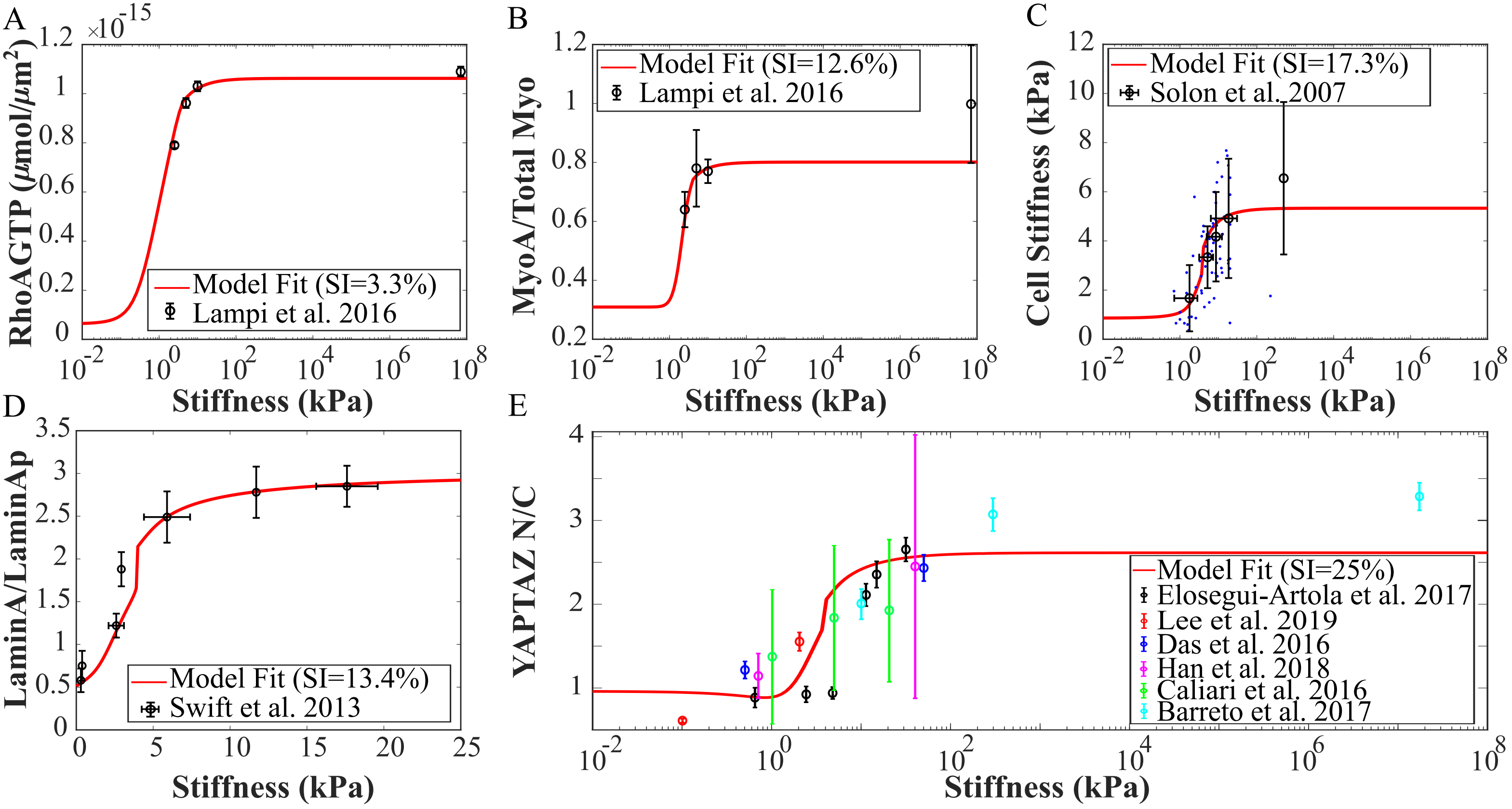
